# Isolation and Characterization of Bacteriocin-Producing Lactic Acid Bacteria from Cheese and Functional Evaluation of Their Synthesized Bioactive Peptides

**DOI:** 10.64898/2026.08.14.744830

**Authors:** Christian Kosisochukwu Anumudu, Taghi Miri, Helen Onyeaka

**Author notes:** Corresponding author email: Christian Anumudu, Helen Onyeaka.

## Abstract

Biopreservatives including nisin and its derivatives are becoming more desirable in the food processing industry because of the growing demand for naturally preserved and minimally processed foods free from artificial preservatives. However, ensuring microbiological safety while meeting these consumer preferences remains a major challenge. This has necessitated the continuous investigation of potential new antimicrobial agents produced by naturally occurring microorganisms. Hence, this study explored the synthesis, characterisation, and optimisation of a bacteriocinogenic lactic acid bacterium and its antimicrobial product, possibly novel bacteriocin (Nisin 2A) from *Lactococcus lactis* isolated from commercial brined cheese. The isolation was achieved by screening for wild-type bacteriocin-producing lactic acid bacteria from dairy products using MRS media. Screening was performed using antagonism assays, yielding five producer organisms. Of these, the isolate whose metabolites exhibited the most potent antimicrobial activity was identified as *Lactococcus lactis*, which synthesised an active antimicrobial peptide designated as Nisin 2A, with a molecular mass of approximately 3.3 kDa as determined by UHPLC-MS and SDS-PAGE. Production of Nisin 2A was scaled up through fed-batch fermentation of *Lactococcus lactis* in modified MRS broth following process optimisation using a Plackett–Burman experimental design and purified by ammonium sulphate precipitation and solid-phase extraction (SPE). Furthermore, the antimicrobial potential of the bacteriocin was evaluated by the agar well diffusion assay and quantified using the tube dilution method. The purified peptide demonstrated broad-spectrum antimicrobial activity, particularly against the test Gram-positive bacteria *Bacillus cereus* and retained its bioactivity across a wide pH range (3–9) and high thermal conditions (up to 100 °C). Furthermore, it had high sensitivity to proteolytic enzymes (Proteinase K and Trypsin). Notably, the peptide was thermostable and retained up to 90% of its initial activity after thermal treatment and maintained consistent inhibitory performance after extended storage. These findings highlight the potential application of Nisin 2A as a natural biopreservative in food systems.

## 1. INTRODUCTION

Modern trends in food processing places emphasis on food safety and the extension of the shelf life of food, while attempting to meet consumers increasing demand for minimally processed foods which are free from chemical preservatives. Given these preferences, along with the growing resistance of pathogens and spoilage bacteria to antimicrobial agents and other chemicals, the food industry is exploring alternative preservation methods (Fair and Tor, 2014). As a result, there is a rising interest in “green technologies,” which include innovative approaches to minimal food processing and the use of microbial metabolites, such as bacteriocins, for the biopreservation of foods on an industrial scale to increase food shelf-life (Deegan et al., 2006, Liu et al., 2022a).

An important biopreservative approach for the control or prevention of the growth of harmful microorganisms in food is the use of lactic acid bacteria (LAB) that produce bacteriocins as starter cultures or the bacteriocin metabolites themselves in food preservation (Coelho et al., 2022, Timothy et al., 2021). LAB are widespread and traditionally have been isolated from various sources, including dairy products, vegetables, and meats (Kariyawasam et al., 2021). These bacteria ferment carbohydrates and produce various organic acids, such as lactic acid, which significantly lowers the pH of fermented foods (Coelho et al., 2022). Similarly, they produce other antimicrobial substances, including hydrogen peroxide, acetaldehyde, and bacteriocins (Setta et al., 2020, Amenu and Bacha, 2023). The Lactic acid bacteria are also known for their antimicrobial activity against both related and other bacterial strains (Zapaśnik et al., 2022, Tang et al., 2022).

Bacteriocins synthesised mainly by LAB are peptides with natural antibacterial properties, often likened to antibiotics, although they are not used for clinical treatments (Darbandi et al., 2022, Meade et al., 2020). They are ribosomally synthesized peptides or proteins that can kill or inhibit the growth of specific target bacteria. Recently, the production of bacteriocins by LAB has been recognized as a probiotic trait, which helps LAB compete within complex microbial communities and positively affects the host’s health (Heilbronner et al., 2021, Canon et al., 2020). They are particularly effective against Gram-positive bacteria, with activity ranging from targeting specific species to multiple species (Tang et al., 2022, Zapaśnik et al., 2022). When applied in foods and food systems, one of the key desirable characteristics is that they do not alter the sensory characteristics of foods, and their use allows for the reduction of other preservative treatments a food receives such as high temperature pasteurization. These characteristics align with modern consumer preferences for minimally processed and more natural food treatments (Knorr and Augustin, 2021, Mesías et al., 2021). Of recent, bacteriocins are preferably added directly to foods as antimicrobial agents, rather than introducing bacteriocin-producing LAB cultures, which could ferment the food’s carbohydrates and change the organoleptic quality of the foods (Zimina et al., 2020). Some examples of bacteriocins tested against spoilage bacteria and pathogens include pediocin and Nisin, enterocin AS-48, bovicin, enterocin 416K1, and bificin C6165. Of these, only Nisin and pediocin have been approved as food additives, with heavy use in the dairy sector while their use in the fruit and vegetable industry remains limited (Bisht et al., 2024, Barbosa et al., 2017). Generally, these bacteriocins are thought to act on bacterial membranes, killing target bacteria by causing membrane permeabilization and extensive pore formation (Kumariya et al., 2019, Pérez-Ramos et al., 2021).

Ongoing research aims to discover newer LAB strains with enhanced properties, such as probiotic potential and bacteriocin production and this constitutes the main focus of this present research as natural products including dairy provide a promising ecological niche for sourcing bacteriocin synthesising LAB. The use of dairy foods as the raw material for this present study is based on both microbiological and cultural reasons. Dairy matrices such as milk, cheese, and fermented milk form one of the richest natural sources of LAB and contain species of *Lactococcus*, *Lactobacillus*, *Streptococcus*, and *Enterococcus*, which are characterized by their high bacteriocinogenic potential (Dapkevicius et al., 2021). Quantitatively, studies have shown that LAB populations in raw or fermented milk products are high at 10 –10⁹ CFU/mL (Ogwaro et al., 2023) and 10⁵–10 CFU/g in fermented vegetable juice (Xu et al., 2018). This richness increases the chances of getting strains with strong antimicrobial activity.

Despite the established use of bacteriocins such as nisin and pediocin in food systems, there remains a need to identify novel bacteriocinogenic LAB strains with enhanced antimicrobial activity and potential application in food biopreservation, particularly from traditional and minimally processed dairy products. Hence, the aim of this study is to examine the diverse native microbiota isolated from dairy products, identify LAB present, synthesize and characterise novel bacteriocins from them. Additionally, the study aims to evaluate the antibacterial activity of these bacteriocin-producing isolates against the foodborne pathogen and spoilage bacteria *Bacillus cereus*.

## 2. METHODOLOGY

### 2.1. Materials

*Bacillus cereus* NCTC 11143 was supplied by the Biochemical Engineering Laboratory, University of Birmingham, UK. Nutrient agar, Mueller–Hinton agar, De Man Rogosa and Sharpe (MRS) broth and agar, Brain Heart Infusion (BHI) broth and agar, Tryptic Soy Broth (TSB), and Maximum Recovery Diluent (MRD) were purchased from Oxoid (Hampshire, UK). Nisin standards (purity ≥ 99%, ≥ 38,000 IU/mg) were obtained from Handary S.A. (Brussels, Belgium). Trifluoroacetic acid (TFA), propan-2-ol, ammonium sulphate, trichloroacetic acid (TCA), acetone, Tris buffer reagents, NaOH, HCl, and other analytical-grade chemicals were purchased from Sigma-Aldrich (UK). Hydrophilic syringe filters (0.22 μm and 0.45 μm) were purchased from Sigma-Aldrich (UK). The API 50 CHL kit for strain characterization was purchased from (BioMérieux, France). Protein quantification was achieved using the BCA Protein Assay Kit (Thermo Fisher Scientific, Rockford, USA) with bovine serum albumin (BSA) standards (Thermo Fisher Scientific, UK). Solid-phase extraction cartridges (C18E-SPE) were sourced from Thermo Fisher Scientific (UK). A BUCHI Rotavapor R-300 rotary evaporator (Fisher Scientific, UK) and a peristaltic vacuum pump (Fisher Scientific, UK) were used. For SDS-PAGE, SDS sample buffer, loading dye and a 1–26 kDa molecular weight marker was purchased from Sigma-Aldrich (UK). A 5 mL HiTrap SP HP cation exchange column was obtained from GE Healthcare (UK). Chromatographic separation and quantification were performed using a Shimadzu Prominence UHPLC system (Japan) equipped with a UV–Photodiode Array Detector and a C18 column (Thermo Fisher Scientific, UK). Mass spectrometry analysis was carried out with a Waters Xevo G2 Q-ToF MS (Waters, Manchester, UK). Enzymes including proteinase K, trypsin, α-amylase, and catalase were purchased from Sigma-Aldrich (UK). Ultrapure water was produced using a Milli-Q Purification System (Merck Millipore, USA). All chemicals and solvents used were of analytical or HPLC grade.

### 2.2. Screening for lactic acid bacteria from dairy sources

Lactic acid bacteria were isolated from food samples (brined cheese) by homogenising the foods in sterile Maximum recovery diluent (MRD) and culturing via the spread plate method on MRS agar. From the resultant colonies, distinct colonies were streaked onto a fresh agar plate to obtain pure cultures according to previously established protocols (Voidarou et al., 2020). These were maintained on agar slants preliminarily and stored at refrigeration temperatures of 4⁰C. From the screening, a total of 38 bacteria isolates were obtained and maintained.

### 2.3. Selected Bacteria Strain and Growth Conditions

*Lactococcus lactis* and LAB 1, LAB 2 and LAB 3 used for this study were isolated from brined cheese as described previously and grown on Brain Heart Infusion (BHI) (Oxoid) media supplemented with 0.5% glucose at 30 ⁰C with shaking at 150 RPM. *Lactobacillus acidophilus* was similarly isolated from brined cheese and grown anaerobically on BHI media at 30 ⁰C while *Bacillus* spp. was grown on nutrient agar (Oxoid) at 37 ⁰C aerobically with shaking at 150 RPM.

### 2.4. Preliminary screening for antimicrobial activity

To test for antimicrobial activity within the growth media, deferred antagonism assay was undertaken by inoculating 10µl of an overnight culture of the producer organism onto the surface of the BHI agar plate by the spread-plate method and incubating for 18 hours. Following colony formation, the resultant colonies were exposed to UV irradiation in a Microbiology Safety Cabinet (MSC chamber) for 1 hour to kill off the colonies. Following the inactivation of the producer organisms, the indicator organism (*Bacillus cereus*) was inoculated into precooled molten nutrient agar. This was used to overlay onto the BHI plate. The diameter of the resultant zones of inhibition was measured using a calliper. This diameter was subtracted from the diameter of the original producer colony and used to calculate the area of the zone of inhibition using the formula α = πr. A total of 5 isolates yielded significant zones of inhibitions and were used for further assays.

### 2.5. Evaluation of the antimicrobial potential of selected isolates

To test the potential of the selected 5 isolates to produce antimicrobial agents, they were inoculated into MRS broth and incubated at 30 ⁰C for 48 hours. Following incubation, every 6 hours, the absorbance of the media at 600nm was measured to estimate the bacteria population. Furthermore, the pH of the media was measured, and an aliquot of the culture was collected and centrifuged at 10,000rpm for 10 minutes. The pH of the supernatant was adjusted to 7 ± 0.2 using 1M NaOH to inactivate any lactic acid produced during growth. This supernatant was subsequently filtered using a hydrophilic 0.22µm filter and utilised for antimicrobial activity testing using the agar well diffusion assay against the indicator organism (*Bacillus cereus*). These well-in-agar assays were incubated at 37 ⁰C for 24 hours and resultant zones of inhibition were measured and the total bacteriocin activity was calculated using the formula.

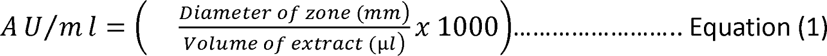

### 2.6. Protein quantification by the BCA assay

Quantification of the protein content of the cell-free extracts was achieved by use of the BCA protein assay kit and Bovine Serum Albumin (BSA) standards as previously described (Lei et al., 2020). Briefly, 0.1ml of the sample was introduced into 2ml of the working reagent containing 50 parts of BCA reagent A with 1 part of BCA reagent B (50:1, Reagent A:B), mixed and incubated at 37 °C for 30 minutes. The absorbance of the preparation was measured using a Jenway spectrophotometer at a wavelength of 562nm. Similarly, a protein calibration curve was obtained using bovine serum albumin (BSA) standards. From obtained BSA standard calibration curve and using a linear regression equation, the concentration of the obtained bacteriocin-like substances was extrapolated.

### 2.7. Preliminary Identification of producer bacteria by the API 50 CHL test assay

The highest producer organisms were preliminarily identified using the API 50 CHL standardised system for evaluating the carbohydrate fermentation profile of lactic acid bacteria (API system, Bio-Merieux, France). This consists of 50 biochemical tests for carbohydrate fermentation. To undertake the assay, 10 ml of distilled water was placed in a sample chamber. Aliquots of the test organism, adjusted to 0.5 MacFarland standard were introduced into the API 50CHL medium (5ml) and were subsequently inoculated into each test well of the strip. The preparation was incubated at 37 ⁰C for 48 hours.

Test results were tabulated as (+/-) based on colour changes in the media and API strips. The results of all biochemical assays were collated and assigned a numeric value as outlined by the manufacturer’s instructions and indicative table. The final species designation was obtained by use of the manufacturers software for identification (apiweb^TM^).

### 2.8. Whole genome sequencing of *Lactococcus lactis*

Producer organisms were grown for 6 hours into the mid-logarithmic phase up to 10 harvested by centrifugation and washed twice using PBS. 500mg of the harvested pellet was transferred into 500ul of the inactivation buffer of DNA/RNA Shield from Zymo Research and transported to the molecular laboratory at -20 ⁰C. Chromosomal DNA was stabilised by the buffer, extracted and sequenced by commercial sequence providers MicrobesNG affiliated to the University of Birmingham using Illumina sequencing. Following sequencing, raw reads were subjected to quality control, de novo genome assembly, and genome annotation by MicrobesNG using their standard bioinformatics pipeline. The assembled genome contigs were provided to the authors for downstream analysis.

The genome of *Lactococcus lactis* was further visualised using the DNA plotter (Carver et al., 2009), whilst the bacteriocin mining tool antiSMASH 7.1.0 (Blin et al., 2024) was employed to identify putative bacteriocin operons within the genome. Furthermore, all the proteins coded for within the genome were aligned against the prokaryotic antimicrobial peptide database with reference to the Protein Basic Local Alignment Search Tool (BLASTP) to obtain a further identification of antimicrobial genes present.

### 2.9. Optimization of bacteriocin production

Bacteria strains were grown using deMann Rogosa and Sharpe (MRS broth). This was augmented with several growth factors (sucrose, inulin, tryptone) for optimized bacteriocin production using a Plackett-Burman (PB) experimental design matrix (Magallanes and Olivieri, 2010). This allowed for the evaluation of the effect of different combinations of the selected growth factors, including environmental factors such as temperature and pH of media on the growth of *Lactococcus lactis* and bacteriocin biosynthesis. *Bacillus cereus* NCTC 11143 was utilized as the indicator organisms. Utilising a two level Plackett-Burman factorial designs allowed for a more rigorous evaluation of the simultaneous effects of these factors at 3 different variable levels on biomass and bacteriocin production. For this assay, 1ml of *Lactococcus lactis* from stock was inoculated into 100ml of MRS broth in an Erlenmeyer flask and incubated at 30 °C for 18 hours with constant shaking at 140rpm. After growth, the bacteria cells were harvested by centrifugation at 5000g for 10 minutes at 4 °C. These were washed twice with PBS to remove all MRS debris and resuspended in Erlenmeyer flasks containing the different growth factors and media according to the experimental design matrix.

### 2.10. Purification of Nisin 2A

*Lactococcus lactis* was grown in modified MRS broth in a 4-litre bench-top fermenter using the optimised media conditions overnight at 30 ⁰C using a fed-batch fermentation approach at an RPM of 150. After 24 hours of growth, the fermentate was centrifuged at 6000 RPM, 4 ⁰C for 30 minutes to separate the supernatant (CFS) from the cells for further purification. Hydrophilic 0.45 μm Millipore syringe filters (Sigma-Aldrich, UK) were utilised for sample filtration. A higher pore size was utilised for large-scale filtration to fast-track the process using a peristaltic Fisherbrand vacuum pump (Fischer Scientific, UK). Also, an additional step to detach potential bacteriocin molecules attached to the separated cells was undertaken by stirring the cell fragments for 4 hours in a mixture of 70% propan-2-ol (IPA) supplemented with 0.1% Trifluoreacetic acid (TFA). This was followed by centrifugation at 6000 RPM for 30 minutes. Following centrifugation, the aqueous/solvent mixture was vaporised to remove the propan-2-ol using a rotary evaporator (BUCHI Rotavapor R-300, Fischer Scientific) set at 42 ⁰C and a pressure of 85mbar, up to 30% of the original volume. Similarly, the supernatant containing the bacteriocin was subjected to ammonium sulphate precipitation by the dropwise addition of ammonium sulphate solution with stirring to the CFS up to a final concentration of 90% w/v at room temperature. Precipitated bacteriocin was harvested by centrifugation at 10,000 RPM for 20 minutes and the supernatant was discarded. The obtained bacteriocin precipitate was dissolved in 200ml of 150ml Tris buffer and 150mM NaCl maintained at a pH of 7.5. Both fractions (from disruption of cell fragments and ammonium sulphate precipitation of the CFS) were combined into one for subsequent purification.

### 2.11. Solid phase extraction (SPE) and SDS-Polyacrylamide Gel Electrophoresis (PAGE)

Following the harvest of the antimicrobial peptide from modified MRS broth as described, a combined protein purification approach was employed as previously described. The extracts were subjected to solid phase extraction (SPE). This was achieved by the use of a C18E-SPE tube. SPE cartridge activation was achieved using 60ml of methanol, followed by 60ml of H_2_O. The aqueous bacteriocin solution was subsequently loaded onto the cartridge. Elution was achieved by the use of 60ml 70% propan-2-ol (IPA) supplemented with 0.1% Trifluoreacetic acid (TFA). Further assays and characterisation of the bacteriocin were undertaken using this crude extract.

Following this, an approximation of the molecular mass of the extract was obtained by SDS–polyacrylamide gel electrophoresis as has been previously described (Schägger and Von Jagow, 1987, Choi et al., 2010), using a 1-26kDa ladder as reference (Sigma-Aldrich, UK). For this assay, 16 μL of the analyte was supplemented with 4 μL 5x SDS sample buffer (0.2 M Tris-HCl, pH 6.8, 10% (w/v) SDS, 40% (v/v) glycerol, 0.02% (w/v) bromophenol blue, and 10 mM dithiothreitol (DTT)). This was loaded onto a tricine gel containing acrylamide. Electrophoresis was run at 100 V for 2 hours. Resultant protein bands were detected via silver staining.

### 2.12. Cation Exchange Chromatography Clean-up of Nisin

Following SPE, analyte was loaded overnight onto a 5 mL HiTrap SP HP cation exchange (cIEX) column (GE Healthcare) with a flow rate of 4 mL/min. Protein elution was monitored by measuring the absorbance at 215 nm as Nisin does not contain any aromatic amino acids and thus cannot be detected at higher wavelengths such as 280 nm. Following the completion of ion exchange chromatography, the column was washed with 50 mM lactic acid at pH 3 to eliminate non-specifically bound material within the column until a uniform baseline was obtained. Bound peptides were eluted initially using 1M NaCl and then by running decreasing concentrations of NaCl through the column at a flow rate of 1 mL/min (optimised at 400mM of NaCl). Following elution, NaCl was removed from the peptides in the elution fractions by precipitating with 20% (v/v) trichloroacetic acid (TCA) overnight at 4’1C. To remove residual TCA, the precipitated protein was washed twice with ice-cold acetone. The purified protein pellet was subsequently suspended in 50 mM lactic acid at pH 3. Concentration of Nisin was determined using the Pierce BCA colourimetric protein assay kit (Thermo scientific) by measuring the absorbance as described by the manufacturers at 584 nm.

### 2.13. Preparative high-performance liquid chromatography-Mass Spectrometry analysis

The obtained eluents were subjected to Liquid chromatography using a Shimazdu Prominence UHPLC system equipped with a UV-Photodiode Array Detector and C18 column (microsob 100-5 Si 250 x 4.6mm) (ThermoFisher Scientific). The mobile phase utilised was A: water with 0.05% TFA and B: a mixture of acetonitrile and methanol acidified with 0.05% Trifluoroacetic acid (TFA). A gradient run was undertaken for 70 minutes wherein the concentration of acetonitrile and methanol was varied from 20% to 75% in 60 minutes. This is followed by a 5% column re-equilibration in ultrapure water. A flow rate of 100µL min-1 and an injection volume of 20 µL was utilised for the HPLC assay. Once retention time was established, the system was realigned into the PrepHPLC format by integrating with the LC-20AP pump for high-flow rate separation. Analyte fraction was collected using a fraction collector for further analysis. Mass analysis was undertaken using a Waters Acquity UPLC equipped with the Xevo-G2 high mass resolution Q-ToF MS/MS instrument using a Waters Acquity UPLC BEH C18 column (2.1 × 50 mm, 1.7 µm) (Waters, Manchester, United Kingdom). The column and the autosampler were maintained, respectively, at +25 °C and +4 °C. The optimization of the mass spectrometry parameters was conducted by direct injection of commercial ready-made Nisin solution standard (Sigma-Aldrich, Dorset, United Kingdom) and assayed using TargetLynx software (Waters Corporation). ESI conditions were set as Ion Spray voltage of 3500 V, Cone voltage: 30 V and Source temperature of 250 °C. The selected reaction monitored (SRM) was optimised with the transitions that rendered the most intensive signal used for quantification of each analyte.

### 2.14. Biochemical Characterisation of Nisin 2A

pH stability of the cell-free extracts was examined by adjusting to 4.0, 6.0, 8.0 and 10.0 using 1M HCl and NaOH. These were incubated at 25 °C for 2 hours and filter sterilised using hydrophilic 0.22 µm syringe filters before their antimicrobial activity was evaluated. Similarly, the cell-free extracts were resuspended in 500ul of PBS (pH 7) and subjected to temperatures of 40, 60, 80 and 100 °C for 30 minutes and allowed to cool to 25 °C before undertaking antimicrobial activity assay (Ndlovu et al., 2015). For enzyme stability assay, 1ml of the cell-free extract was mixed with 100 μl of prepared enzyme solution (Proteinase K, trypsin, α-amylase and catalase) and incubated for one hour at 37 °C before undertaking antimicrobial activity assay against the test organism by the agar well diffusion assay (Ndlovu et al., 2015). The resulting areas of inhibition were measured. pH buffers and untreated samples (temperature and enzymes) were used as controls.

All experiments were done in triplicates.

## 3. RESULTS AND DISCUSSIONS

### 3.1. Biochemical Identification of Isolates

In this study, the API 50 CHL system (Figure 1) was employed for the preliminary identification of five of the most promising LAB isolates based on their carbohydrate fermentation profiles with the aid of the apiweb software. The most active isolate was identified as *Lactococcus lactis*, followed by Lactobacillus acidophilus, with other isolates including Lactobacillus brevis, Lactobacillus fermentum, and Pediococcus pentosaceus (Table 1).

**Figure 1.**
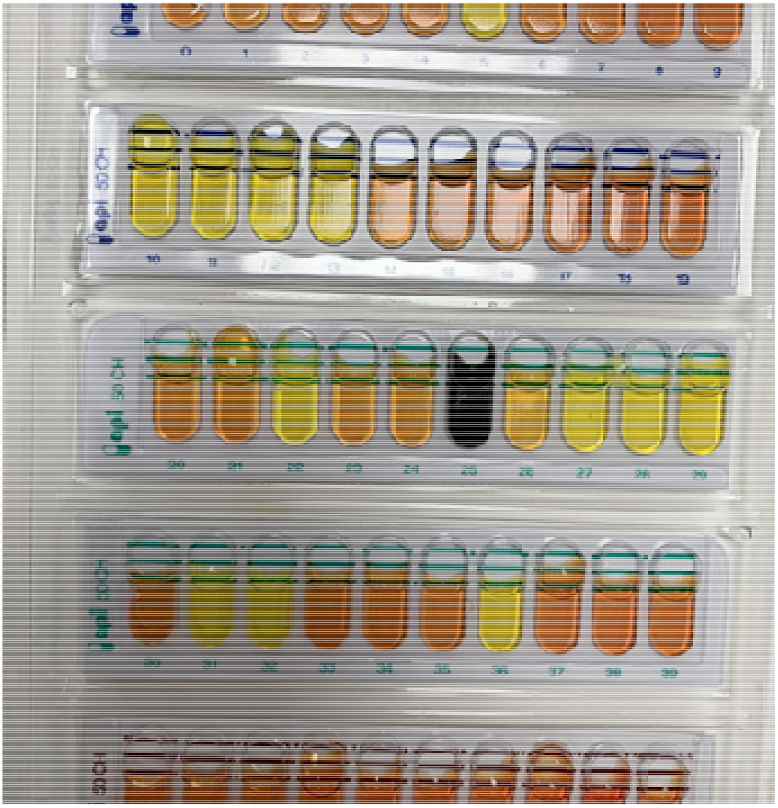
API 50 CHL standardized assay for LAB identification. The API 50 CHL test kit is a widely validated phenotypic system that profiles bacterial fermentation of 49 carbohydrates and esculin (Jeyagowri et al., 2023). Isolate 1 (*Lactococcus lactis*) demonstrated broad fermentative capability and produced the highest antimicrobial activity. This aligns with prior studies showing that *L. lactis* efficiently ferments multiple sugars, including glucose, lactose, and maltose, and produces Nisin, a well-characterized bacteriocin (Kondrotiene et al., 2023). Isolate 2 (*Lactobacillus acidophilus*) which is a well-established probiotic genera, showed moderate antimicrobial activity and a carbohydrate utilization pattern consistent with heterofermentative metabolism (Bian, 2008). *Lactobacillus brevis, L. fermentum, and Pediococcus pentosaceus* showed limited or selective sugar fermentation, which is consistent with their more specialized ecological roles (Bintsis, 2018).

**Table 1.** Identification of producer organisms using the API 50 CHL standardised system.

| Isolate | Organism |
| --- | --- |
| Isolate 1 | *Lactococcus lactis* |
| Isolate 2 | *Lactobacillus acidophilus* |
| Isolate 3 | *Lactobacillus brevis* |
| Isolate 4 | *Pediococcus pentosaceus* |
| Isolate 5 | *Lactobacillus fermentum* |

### 3.2. Antimicrobial potential of isolates

Selected lactic acid bacteria were screened for antimicrobial activity against *B. cereus.* One of the isolates (*Lactobacillus acidophilus*) had limited activity against the test organism while another isolate (*Lactococcus lactis*) had a more pronounced antimicrobial activity, the other three producer organisms (*Lactobacillus brevis, Pediococcus pentosaceus* and *Lactobacillus fermentum*) had intermediate activity (Table 2). The zone of inhibition of *L. lactis* increased significantly from 6.6 ± 0.014 mm at 6 hours incubation time to 15.1 ± 0.016 mm at 30 hours, suggesting that this organism produces a potent antimicrobial compound, potentially a bacteriocin, which is very effective against *B. cereus* during its peak growth phase. The coincidental drop in pH of the media to 4.2 ± 0.011 during the highest activity of *L. lactis* (15.1 ± 0.016 mm zone of inhibition) confirms the production of organic acids which have been known to synergistically enhance the activity of bacteriocins. Notably, the antimicrobial activity of *L. lactis* declined slightly after the 30-hour incubation period to 13.6 ± 0.015 mm at 48 hours, which may have been because of depletion of nutrients in the media, or feedback inhibition that usually occurs to prevent the overproduction of metabolites that may not be needed anymore (Lorendeau et al., 2015). The other organisms showed limited activity against the test organism. While *L. acidophilus* showed good antimicrobial activity with a maximum zone of inhibition of 13.0 ± 0.015 mm at 24 hours, *L. brevis, P. pentasaceus,* and *L. fermentum* showed moderate antimicrobial activities, with maximum zones of inhibition ranging from 8.0 mm to 9.1 mm. The observed difference in the antimicrobial activity among these organisms may be because of the difference in their abilities to produce bacteriocins, secrete organic acid, or metabolic efficiency. The high antimicrobial activity of *L. lactis* as observed in this study is in agreement with the observations made by Akbar et al. (Akbar et al., 2019) where the *L. lactis* subsp. *lactis* isolated from fermented milk in their study showed antibacterial activity against notable foodborne pathogens used as target bacteria including *Salmonella typhimurium* (11.2 ± 1.72 mm)*, Staphylococcus aureus* (21.2 ± 1.2 mm)*, Escherichia coli* (13.4 ± 1.15 mm) and *Listeria monocytogenes* (27.3 ± 1.4 mm). The enhanced antimicrobial efficacy due to the synergistic effect of organic acids in combination with bacteriocins observed in this study as shown by the gradual drop in pH and increase in antimicrobial activity correlates with the findings of Zhang et al. (Zhang et al., 2019) and Soltani et al. (Soltani et al., 2022). These demonstrate that *L. lactis* has appreciable potential as a biopreservative in the food industry.

**Table 2.** Growth of producer organism and antibacterial activity of cell free extract.

| Incubation Time (h) | Parameter | Isolate 1: *L. lactis*<br>(Mean ± SD) | Isolate 2: *L. acidophilus* (Mean ± SD) | Isolate 3: *L. brevis*<br>(Mean ± SD, SE) | Isolate 4: *P. pentosaceus* (Mean ± SD) | Isolate 5: *L. fermentum* (Mean ± SD) |
| --- | --- | --- | --- | --- | --- | --- |
| 6 | Biomass (Abs<sub>600</sub>) | 0.34 ± 0.013 | 0.30 ± 0.011 | 0.32 ± 0.012 | 0.28 ± 0.013 | 0.30 ± 0.012 |
| | Avg ZOI (mm) | 6.6 ± 0.014 | 5.1 ± 0.013 | 4.1 ± 0.012 | 4.0 ± 0.012 | 4.3 ± 0.013 |
| | pH of media | 6.4 ± 0.011 | 6.4 ± 0.012 | 6.2 ± 0.011 | 6.3 ± 0.013 | 6.2 ± 0.012 |
| 12 | Biomass (Abs<sub>600</sub>) | 0.47 ± 0.014 | 0.46 ± 0.013 | 0.50 ± 0.015 | 0.44 ± 0.013 | 0.52 ± 0.015 |
| | Avg ZOI (mm) | 8.9 ± 0.013 | 7.3 ± 0.014 | 5.4 ± 0.012 | 5.4 ± 0.013 | 4.8 ± 0.013 |
| | pH of media | 6.2 ± 0.011 | 6.0 ± 0.012 | 6.0 ± 0.011 | 6.2 ± 0.013 | 6.2 ± 0.012 |
| 18 | Biomass (Abs<sub>600</sub>) | 0.86 ± 0.015 | 0.90 ± 0.014 | 0.84 ± 0.013 | 0.78 ± 0.014 | 0.82 ± 0.013 |
| | Avg ZOI (mm) | 11.2 ± 0.015 | 11.2 ± 0.014 | 8.2 ± 0.013 | 8.0 ± 0.013 | 7.8 ± 0.012 |
| | pH of media | 5.0 ± 0.012 | 4.8 ± 0.011 | 4.6 ± 0.011 | 4.6 ± 0.012 | 4.4 ± 0.012 |
| 24 | Biomass (Abs<sub>600</sub>) | 1.64 ± 0.016 | 1.10 ± 0.013 | 1.04 ± 0.013 | 0.98 ± 0.014 | 1.12 ± 0.015 |
| | Avg ZOI (mm) | 15.0 ± 0.016 | 13.0 ± 0.015 | 8.4 ± 0.014 | 8.2 ± 0.013 | 7.6 ± 0.013 |
| | pH of media | 4.0 ± 0.011 | 4.2 ± 0.012 | 4.0 ± 0.011 | 3.9 ± 0.012 | 4.2 ± 0.012 |
| 30 | Biomass (Abs<sub>600</sub>) | 2.00 ± 0.015 | 1.82 ± 0.015 | 1.60 ± 0.014 | 1.35 ± 0.013 | 1.48 ± 0.013 |
| | Avg ZOI (mm) | 15.1 ± 0.016 | 12.0 ± 0.015 | 9.1 ± 0.014 | 7.8 ± 0.013 | 8.0 ± 0.013 |
| | pH of media | 4.2 ± 0.011 | 4.0 ± 0.011 | 3.8 ± 0.011 | 3.4 ± 0.010 | 4.0 ± 0.011 |
| 36 | Biomass (Abs<sub>600</sub>) | 2.32 ± 0.016 | 2.10 ± 0.015 | 2.30 ± 0.015 | 1.90 ± 0.014 | 2.22 ± 0.014 |
| | Avg ZOI (mm) | $14.2 \pm 0.015$ | $11.9 \pm 0.014$ | $8.8 \pm 0.013$ | $8.0 \pm 0.013$ | $8.0 \pm 0.013$ |
| | pH of media | $3.5 \pm 0.011$ | $3.8 \pm 0.011$ | $3.7 \pm 0.011$ | $3.3 \pm 0.010$ | $3.8 \pm 0.011$ |
| **42** | Biomass ($\text{Abs}_{600}$) | $2.36 \pm 0.016$ | $2.20 \pm 0.015$ | $2.25 \pm 0.015$ | $2.25 \pm 0.015$ | $2.56 \pm 0.016$ |
| | Avg ZOI (mm) | $14.3 \pm 0.015$ | $11.3 \pm 0.014$ | $9.0 \pm 0.013$ | $7.8 \pm 0.013$ | $7.2 \pm 0.012$ |
| | pH of media | $3.7 \pm 0.011$ | $3.7 \pm 0.011$ | $3.5 \pm 0.010$ | $3.6 \pm 0.010$ | $3.6 \pm 0.011$ |
| **48** | Biomass ($\text{Abs}_{600}$) | $2.34 \pm 0.016$ | $2.20 \pm 0.015$ | $2.50 \pm 0.016$ | $2.34 \pm 0.016$ | $2.54 \pm 0.016$ |
| | Avg ZOI (mm) | $13.6 \pm 0.015$ | $11.1 \pm 0.014$ | $8.8 \pm 0.013$ | $7.8 \pm 0.013$ | $7.4 \pm 0.013$ |
| | pH of media | $3.3 \pm 0.010$ | $3.6 \pm 0.011$ | $3.5 \pm 0.010$ | $3.6 \pm 0.010$ | $3.4 \pm 0.010$ |

As outlined in Table 3, of the 5 isolates tested, *L. lactis* demonstrated the highest antimicrobial activity, with the largest zone of inhibition (15.1 ± 0.012 mm), the highest total protein content of 54.2 mg/L (as measured using the BCA assay). For quantification of antimicrobial activity, the tube dilution assay was undertaken to determine the minimum inhibitory concentration (MIC). Also, Table 3 presents the antimicrobial characteristics of the cell free extracts from the isolates as determined by the tube dilution assay, highlighting the MIC. Similar to the results obtained in the agar-well diffusion assay, extracts from *L. lactis* had the lowest MIC (50 µg/ml), in contrast to the other organisms that demonstrated lower antimicrobial efficacy, smaller zones of inhibition, lower protein content and higher MIC values. Due to the potency of the synthesised antimicrobial peptides of *L. lactis* in this study, it was selected for further examination.

**Table 3.**
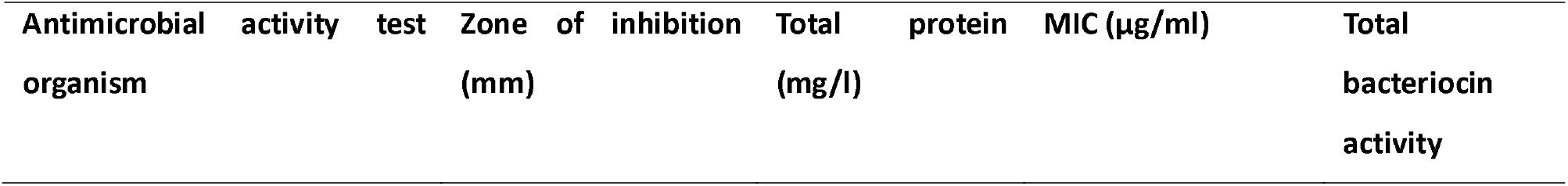
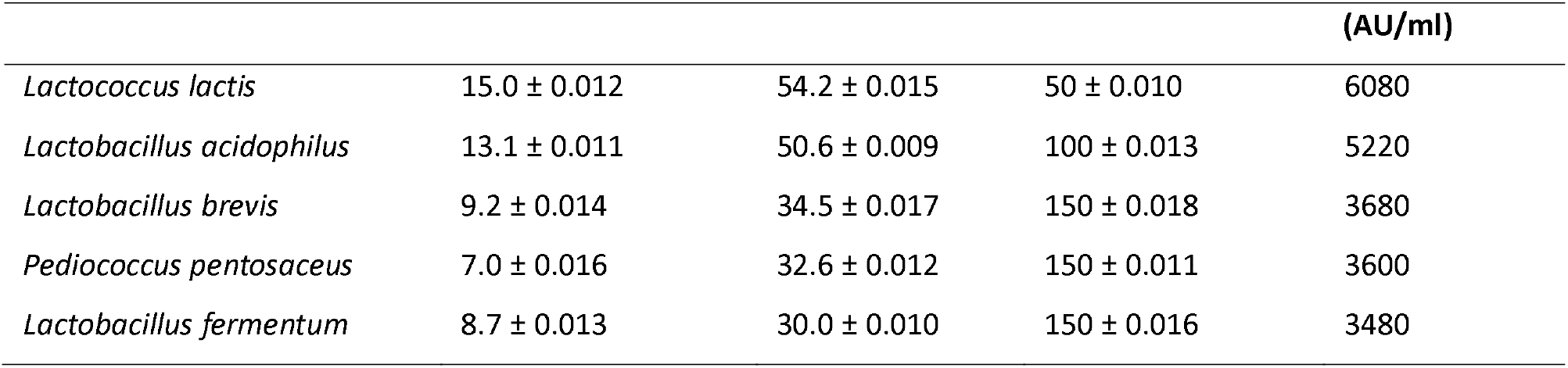
MIC of the cell free extract from the isolates.

The high protein content in the cell-free extract of *L. lactis* observed in this study correlates with its strong antimicrobial activity, in contrast to the other isolates that produced lower protein concentrations and exhibited weaker antimicrobial activity, suggests that the bacteriocins produced by *L. lactis* which are proteinaceous in nature, are possibly the major contributors to its inhibitory effect against the test bacterium (B. cereus) (Anumudu et al., 2021). The lower MIC values of *L. lactis* further supports this observation about its potency, since only a smaller concentration of its extract was required to inhibit the growth of the test organism. A previous study by Lay et al. (Lay et al., 2016) demonstrated the inhibitory effects of Nisin produced by several strains of *L. lactis* against clinical isolates of Clostridium difficile at low concentrations (6.2 μg ml for Nisin Z and 0.8 μg ml for Nisin A), which is consistent with the low MIC and high antimicrobial activity observed in this study. Also, as suggested in another study by Abbasiliasi et al. (Abbasiliasi et al., 2017), the variation in the total bacteriocin activity and antimicrobial activity among the strains of different LAB species isolated in this study demonstrates the difference in their ability to produce bacteriocins, which can be ascribed to several reasons including genetic factors and environmental conditions.

The highest activity was recorded by the extract from *Lactococcus lactis*. This organism was selected and utilised for further studies due to the potency of the synthesised antimicrobial peptides. Importantly, the producer organism *Lactococcus lactis* is an important starter organism in many fermentation processes (Jang et al., 2015) and has been shown to be a source of several enzymes and bacteriocins (Akçelik et al., 2006).

### 3.3. Whole Genome Sequencing and Identification of the putative gene cluster responsible for bacteriocin synthesis in *Lactococcus lactis*

16S rRNA gene sequencing is considered one of the most efficient tools in the generic and specific identification of isolates as the 16S rRNA gene possesses conservative properties which could provide sufficient resolution for identification (Yu et al., 2017). With the advances made in sequencing and computational technologies, it is now possible to provide accurate identification of organisms through the combination of core genome phylogeny and taxonomic guidance based on whole-genome sequences (Xu Jianping, 2006). Whole genome sequencing and gene annotation were undertaken to first identify the organism and then to possibly map the biosynthetic genes which are responsible for the antimicrobial activity of the *Lactococcus lactis*. Genome annotation was undertaken using the RAST tool kit (RASTtk) and a unique genome identifier of 1358.2762 was assigned. The taxonomy of the organism was identified as follows;

Bacteria> Terrabacteria group> Bacillota> Bacilli> Lactobacillales> Streptococcaceae> *Lactococcus*> *Lactococcus lactis*

The Pathosystems Resource Integration Center (PATRIC) which is a comprehensive bioinformatics resource that is focused on bacterial genomes was utilised for phylogenetic analysis by comparing the genome to the database maintained by the National Centre for Biotechnology Information (NCBI) (Wattam et al., 2017). Using PATRIC, a comprehensive genome analysis report was obtained, and the protein sequences present within the families were aligned, with the nucleotides for each of the sequences mapped to the protein. The phylogenetic tree is shown in Figure 2. The phylogenetic result showed that the *Lactococcus lactis* strain isolated in this study was closely related to other species of the same genus (Lactococcus). The phylogenetic relationship across the *Lactococcus* species is supported by 100% of the bootstrap iterations. Two branches were clearly identified with a total of 12 species including the reference strain. The first branch contained 3 *Lactococcus* subspecies which includes *L. lactis* III403272623, *L. lactis* lac001 1358.2 Strain762 1358.2762 (reference), *L. lactis* subsp. Cremoris TIFN6 1234876.3 while the second one contained S. thermophilus M17PTZA496 1433289.7, S. thermophilus strain S9 1308.54, S. thermophilus CNCM1-1630 1042404.3 and S. pseudopneumoniae ST7493 10544460.4. The inclusion of Streptococcus and Enterococcus species in the phylogenetic analysis served as closely related comparative taxa within the lactic acid bacteria group, allowing for clearer resolution of inter-genus relationships and supporting the placement of the study isolate within the *Lactococcus* clade. The result suggests that the reference strain L. Lactis lac001 1358.2762 1358.2762 is most closely related to *L. lactis* III403272623 and, *L. lactis* subsp. Cremoris. The *Lactococcus lactis* total genome size was 2522937 bp. A smaller genome size was suggested to indicate adaptations for reproductive efficiency or competitiveness in new environments (Burke and Moran, 2011). Such genome streamlining is consistent with adaptation to nutrient-rich dairy environments, where metabolic efficiency and rapid growth confer a selective advantage to lactic acid bacteria (Sun et al., 2025).

**Figure 2.**
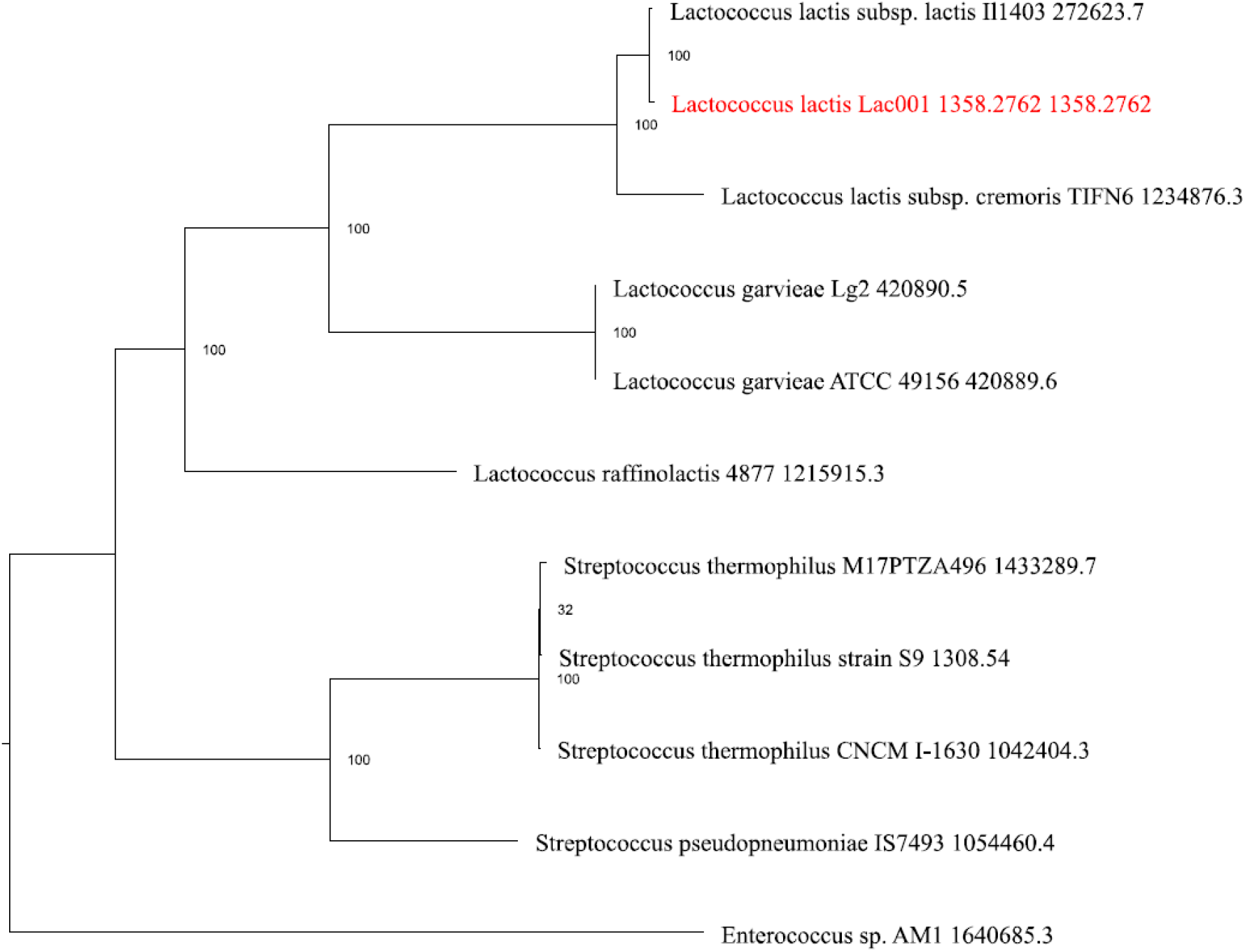
Phylogenetic tree showing 100% genetic relatedness between *Lactococcus lactis* lac001 1358.2762 1358.2762 and other strains of *Lactococcus species* (*L. lactis* III403272623 and *L. lactis* subsp. *Cremoris*)

Table 4. shows the protein features of *L. lactis*, with about 650 proteins associated with enzymatic functions, which suggests that this *L. lactis* strain has a wide range of metabolic capabilities, including possible biosynthesis of antimicrobial compounds like bacteriocins and organic acids. Similar studies on genome sequencing of *L. lactis* such as the one conducted by Mileriene et al. (Mileriene et al., 2023) have shown that functional assignments of proteins often include pathways for the biosynthesis of antimicrobial compounds which corroborates the findings in this present study. While 2,024 proteins known for different functional assignments were found, the 624 hypothetical proteins that were present indicate that some aspects of this organism’s genome are still poorly understood. Similarly, it had 566 proteins with Gene Ontology (GO) assignments, and 474 proteins that were mapped to KEGG pathways. PATRIC annotation includes two types of protein families, and this genome has 2538 proteins that belong to the genus-specific protein families (PLFams) for, and 2567 proteins that belong to the cross-genus protein families (PGFams). Hence, more functional studies are required on these hypothetical proteins as they may reveal additional novel antimicrobial agents or traits that contribute to their stress response capabilities.

**Table 4.** Protein features of *Lactococcus lactis*.

| Proteins | Numbers |
| --- | --- |
| Hypothetical proteins | 624 |
| Proteins with functional assignments | 2,024 |
| Proteins with EC number assignments | 650 |
| Proteins with GO assignments | 566 |
| Proteins with Pathway assignments | 474 |
| Proteins with PATRIC genus-specific family (PLfam) assignments | 2,538 |
| Proteins with PATRIC cross-genus family (PGfam) assignments | 2,567 |

### 3.4. Genomic Annotation of *Lactococcus lactis*

Many of the annotated genes are homologous to known transporters, virulence factors, drug targets, and antibiotic resistance genes. The number of genes and the specific source database are provided in Appendix 3.1 and 3.2. PATRIC annotation of genes is presented in Figure 3.

**Figure 3:**
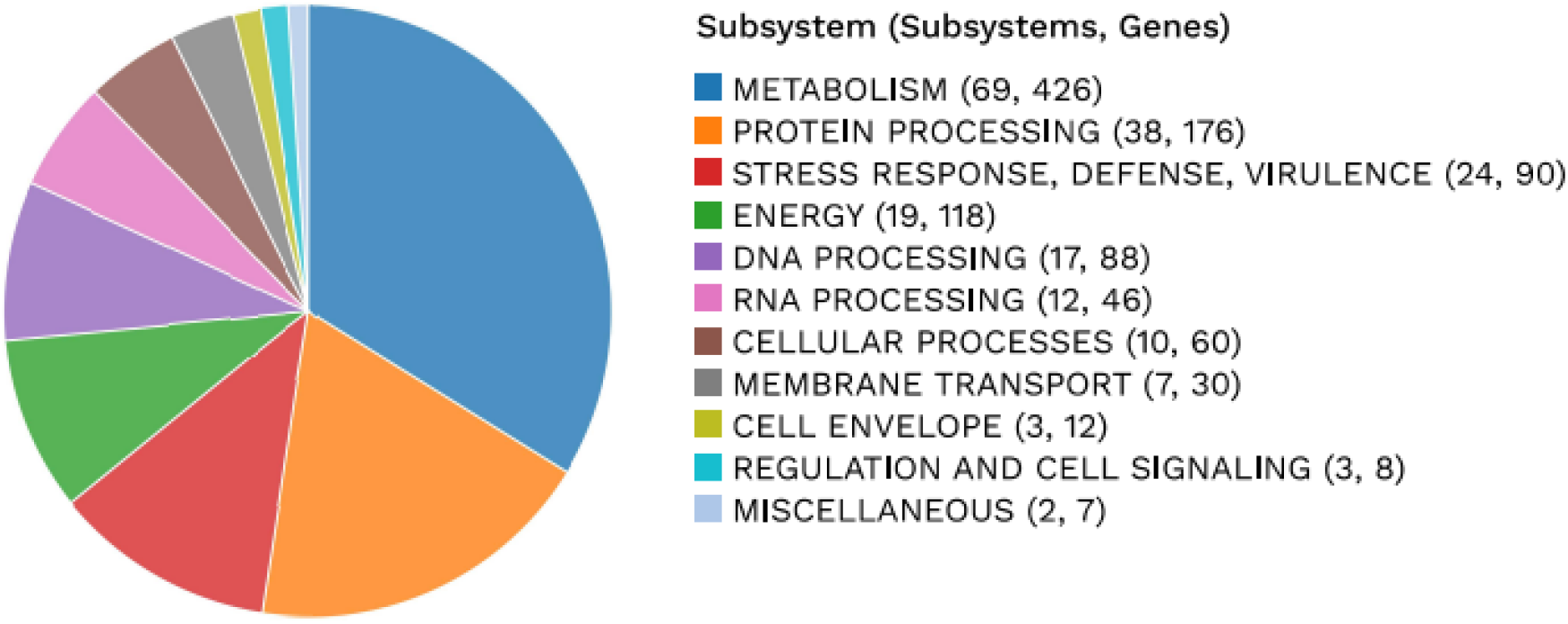
PATRIC annotation analysis of the subsystems unique to *L. Lactis* Lac001 genome. The subsystem distribution of *Lactococcus lactis* reveals that metabolism represents the largest proportion of annotated genes, with 69 subsystems and 426 genes. This dominance of metabolism-associated genes underscores the central role of carbohydrate and energy metabolism in *L. lactis*, reflecting its ecological adaptation to nutrient-rich dairy environments such as milk, where efficient carbohydrate utilization and energy extraction are essential for growth and fermentation activity (Kleerebezem et al., 2020). This metabolic bias is consistent with previous reports showing that most dairy isolates of *L. lactis* exhibit a relatively limited capacity to utilize a broad range of carbohydrates, a feature that has been attributed to genome decay associated with prolonged cultivation in nutrient-rich niches such as milk (Wels et al., 2019). Such reductive evolution favors the retention and optimization of core metabolic pathways required for growth in dairy matrices, rather than metabolic versatility. Similar patterns of metabolic streamlining and specialization have been observed in other dairy-associated lactic acid bacteria, including *Streptococcus thermophilus* and *Lactobacillus helveticus* (Kleerebezem et al., 2020), suggesting that the metabolic dominance observed in *L. lactis* reflects a broader evolutionary trend among LAB adapted to dairy fermentation environments.

Furthermore, protein processing was observed to be the second most abundant category, with 38 subsystems and 176 genes. A high proportion of protein processing genes supports the capacity of the organism for active protein synthesis and post-translational modification, processes that are necessary for the production and secretion of peptides, including antimicrobial peptides such as bacteriocins (Kondrotiene et al., 2023). In lactic acid bacteria, efficient protein processing also contributes to stress tolerance and rapid adaptation during fermentation. This is reflected in the prominence of protein processing subsystems in the isolated *Lactococcus lactis*, which include genes involved in proteolysis, peptide transport, and protein quality control. The proteolytic and protein quality control systems of LAB are central to both growth and stress resilience as they enable the breakdown of extracellular proteins into peptides and amino acids that fuel metabolism and support cellular functions (De Angelis et al., 2016). Such systems have been shown to support cellular performance under fermentation-associated stresses, including acidification and nutrient fluctuations commonly encountered in dairy environments (Wang et al., 2021).

Figure 4. presents a circular graphical display showing the distribution of the genome. The image depicts from outer to inner rings, the contigs, CDS on the forward strand, CDS on the reverse strand, RNA genes, CDS with homology to known antimicrobial resistance genes, CDS with homology to know virulence factors, GC content and GC skew. The colours of the CDS on the forward and reverse strand indicate the subsystem that these genes belong.

**Figure 4.**
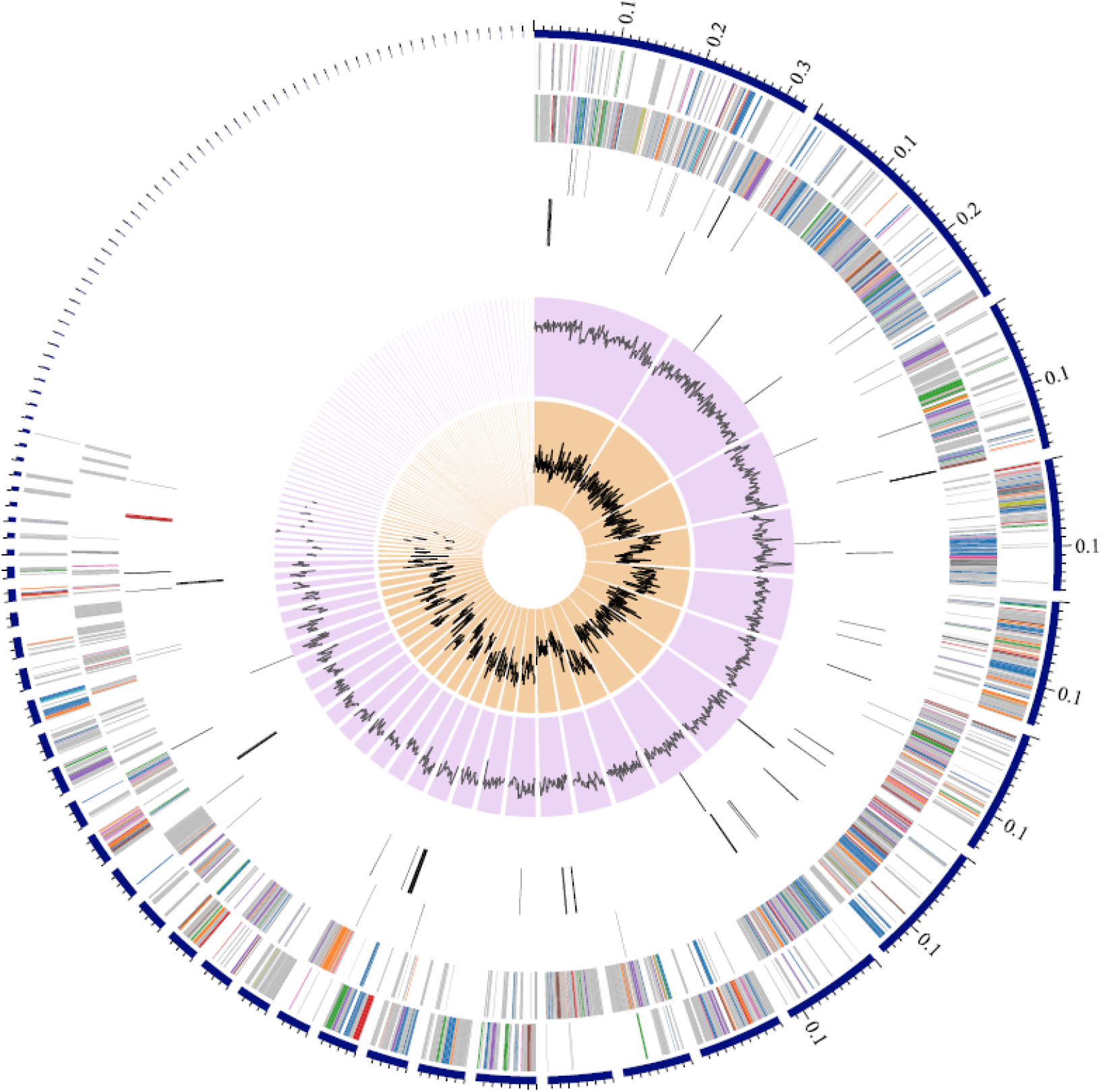
A circular graphical display of the distribution of the genome annotations Following whole genome sequencing, the genome was screened using antiSMASH 7.1.0 web tool to identify the putative gene clusters encoding for antimicrobial activity. Both analyses predicted the presence of highly hydrophobic circular bacteriocin with an amino acid sequence similar to Nisin as shown in Figure 5.

Bacterial genes code for the metabolism of various compounds resulting in a diversity of bioactive compounds. Some of these coded bioactive compounds are of potential pharmaceutical value including antibiotics, cholesterol-lowering drugs, and antitumor drugs. Here, antiSMASH 7.1.0 was used to predict metabolic pathways (Blin et al., 2024). Data mining of genomic and metagenomic sequences was utilised as an important strategy for identifying bacteriocin producers. This is a promising approach because many features of bacteriocin gene clusters, and especially bacteriocin modification genes are highly conserved (Oliveira et al., 2017). As shown in Figures 5 and 6, regions of five different secondary metabolite biosynthetic gene clusters (BGCs) were detected in the *L. lactis* genome which are ribosomally synthesized and post-translationally modified peptide (RiPP-like), RiPPs produced by radical SAM (S-adenosylmethionine) enzyme (RaS-RiPP), type III polyketide synthases (T3PKS), Lanthipeptide-Class-I, and Betalactone. These regions include the most similar known clusters of gene that code for Nisin A (Figure 5). The presence of loci encoding RiPP-like metabolites such as lactococcin indicate the ability of the strain to produce ribosomally synthesized peptides with bactericidal or bacteriostatic effect, which are critical for its role in food preservation (Zhang et al., 2022). Lactococcin is a class II bacteriocin with a molecular weight of 5 KDa which majorly inhibits the growth of sensitive lactococci, although its bactericidal ability depends on the reduced state of Cys-24 residue (Mileriene et al., 2023). With the ability of lactococcin to target and lyse specific bacterial strains, including other *Lactococcus* strains used in starter cultures, their producer strains have potential applications in the dairy industry, as the intracellular enzymes produced during their controlled lysis can aid in the breakdown of proteins and fats, thereby accelerating the ripening process and increasing flavour development (Mileriene et al., 2023, Zhang et al., 2022). Different types including lactococcin A, B, and M, have been confirmed in other strains of *L. lactis* subsp. lactis isolated in the study by Pisano et al. (Pisano et al., 2015).

**Figure 5.**
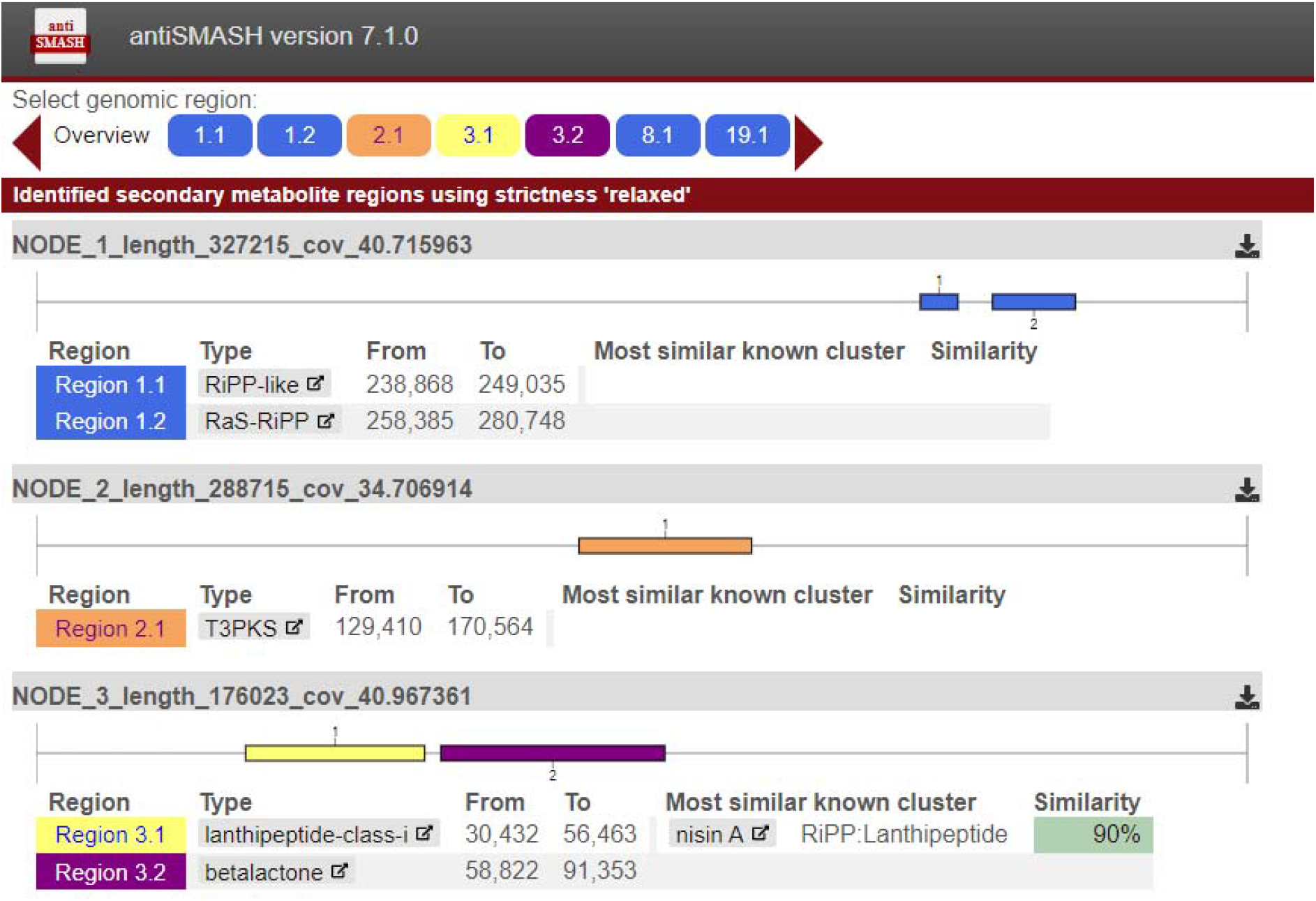
Linear map of biosynthetic gene clusters which code for putative bacteriocin Lantipeptide class I, beta lactone T3PKS and Nisin having a 90% similarity.

**Figure 6.**
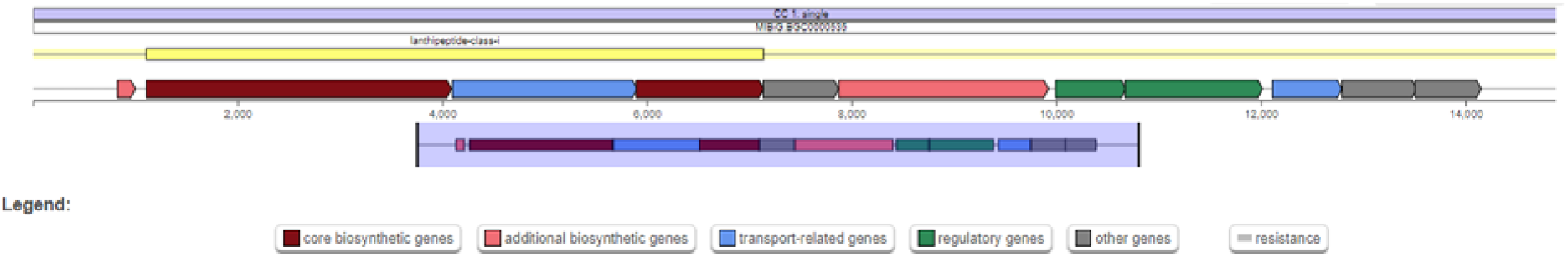
Metabolite-producing regions RiPP-like (A), RaS-RiPP (B), T3PKS (C), Lanthipeptide-Class-I (D), and Betalactone (E) in the genome of *L. lactis* lac001 1358.2762 1358.2762 detected with the antiSMASH 7.1.0 web tool. The arrows indicate the direction of transcription for each gene and the colours indicate gene function including biosynthesis, transport, and regulation.

RaS-RiPP BGCs have been found to be responsible for the biosynthesis of RiPPs, and typically include genes that encode a precursor peptide, a RaS enzyme, and additional genes for transport and specific modifications (Benjdia et al., 2017). In the genome of *L. lactis* Lac001, three distinct regions of Ras-RiPP were identified suggesting a greater diversity of RiPP production in the strain, which potentially enhances its functional versatility. A similar study by Tenea et al. (Tenea, 2023) detected two Ras-RiPP BGCs in the *L. lactis* strains isolated in their study, suggesting that further studies were needed to fully assess the impact of these regions on the biotechnological potential of the strain. T3PKS is one of the two most abundant BGCs in all LAB genera, and are known to produce bacteriocins involved in food safety (Mileriene et al., 2023). The presence of T3PKS in *L. lactis* Lac001 genome producing enzymes such as hydroxymethylglutaryl-CoA synthase, increases the metabolic potential of the strain to encompass polyketide biosynthesis, a broad family of secondary metabolites with a variety of biological activity and potential therapeutic uses (Josephs-Spaulding et al., 2024). The study by Okoye et al. (Okoye et al., 2022) found that T3PKS was present in the genomes of five LAB species isolated from fermented vegetables including *Lactobacillus plantarum, Pediococcus pentosaceus, Weissella hellenica, Lactobacillus buchneri*, and *Enterococcus* sp., fortified with distinct genes that encode hydroxymethylglutaryl-CoA synthase, which has the ability to demonstrate antibacterial action and biopreservative potential.

Lanthipeptides are a subclass of RiPPs, produced by a wide variety of microorganisms including various strains of LAB, and are characterized by the presence of lanthionine and methyllanthionine residues, which are formed through modifications of serine, threonine, and cysteine residues by enzymes (Fu et al., 2023). Lanthipeptide-class-I BGCs encode the enzymes and regulatory proteins required for the synthesis, modification, and transport of these peptides. Its presence in *L. lactis* Lac001 genome encoding enzymes from lanthionine synthetase C family for instance, suggests the ability of this strain to produce lantibiotics, a class of antimicrobial peptides with broad-spectrum activity against competing microorganisms, including resistant bacterial pathogens (Garvey, 2023). *L. lactis* strains isolated from traditional ewe’s and goat’s raw milk cheeses in the study by Garcia-Cayuela et al (García-Cayuela et al., 2017), were found to produce lantibiotics including Nisin and lacticin. The betalactone BGCs detected in *L. lactis* Lac001 genome, encoding biosynthetic genes such as acetyl-CoA carboxylase biotin carboxylase and transport-related genes which collectively contribute to the production and transportation of betalactone compounds (Presented in Figure 6), emphasizes the potential of this strain for food preservation (Du et al., 2023). Other secondary metabolites of the biosynthetic gene clusters in *Lactococcus lactis* is provided in Appendix 3.3.

In this study, full-length sequencing of the 16S rRNA gene allowed accurate identification of the isolate as *Lactococcus lactis*. Further visualization of the genome using the DNAPlotter tool (Carver et al., 2009) enabled a structural overview of the genome architecture, while phylogenetic analysis revealed that the identified isolate (*L. lactis* lac001 1358.2762) shares the closest evolutionary relationship with *L. lactis* strain III403272623 and *L. lactis* subsp. *cremoris*. These findings underscore the resolution power of phylogenomic analysis in delineating strain-level relationships within LAB species.

The whole-genome sequencing of *Lactococcus lactis* revealed a total genome size of 2,522,937 base pairs, assembled into 99 contigs, with a GC content of 34.92%. These values are similar to those reported by other studies (Mileriene et al., 2023, Liu et al., 2022b) for genomes of other *L. lactis* strains isolated from dairy products. According to Bobay and Ochman (Bobay and Ochman, 2017), reduced genome sizes may reflect bacterial adaptation toward increased reproductive efficiency or specialization to specific environmental niches. The moderate GC content is consistent with that typically reported for LAB and reflects evolutionary influences such as environmental factors, mutation bias, host interaction, and ecological adaptation (Wu et al., 2012). Through the analysis of the trends in pan-genome size of *Lactococcus* species/subspecies it is shown that *Lactococcus* genus has an open pan-genome. The presence of this may be associated with the diverse range of environments *Lactococcus* species can colonize and the existence of numerous ways of exchanging genetic material. It has been shown by previous studies that bacterial genomes change when they adapt to variable conditions, and for greater niche diversity, larger pan-genomes are required (Konstantinidis and Tiedje, 2004). However, no plasmids were found in the *L. lactis* strain isolated in this study which contradicts previous findings (Liu et al., 2022b, Mileriene et al., 2023). Although plasmids of *L. lactis* have been found to be associated with a range of essential functions including bacteriocin production, utilization of citrate, and hydrolysis of proteins, they have also been found to harbour genes that confer resistance to antibiotics (Cui et al., 2015, Mileriene et al., 2023). Its absence in *L. lactis* Lac001 could be because of the absence of selective pressures that promote the retention of plasmids, the environmental niche the strain was isolated from, or its evolutionary history (Rodríguez-Beltrán et al., 2021).

Although there was no success in the identification of the specific putative genes responsible for coding Nisin 2A in this study using antiSMASH 7.1.0 (Blin et al., 2024), this is not of much concern as this may be due to a lack of information on this variant of Nisin produced by the *Lactococcus lactis* or a higher level of diversity compared to previously mapped bacteriocins from this organism. This requires further investigation as numerous studies have utilised genomic data for in silico bacteriocin identification including; formicin (Collins et al., 2016), and novel bacteriocins from *Lactobacillus rhamnosus* L156.4 strain (Oliveira et al., 2017). Overall, as the complete sequence of the genes responsible for the biosynthesis of the bacteriocin is not yet known, it is not possible to establish the true novelty or otherwise of the synthesized antimicrobial peptide. This can be explored in subsequent studies as the discovery of novel antimicrobial compounds especially with potential application in the food sector is important currently, with the surge in antimicrobial resistance and the drive in naturally preserving foods using biomolecules.

### 3.5. Optimization of bacteriocin production by the Plackett-Burman experimental design

To evaluate the simultaneous effect of supplementation of growth medium (MRS) on the growth of *Lactococcus lactis* and bacteriocin production, Plackett-Burman experimental design as shown in Table 5 was utilized to change the growth conditions of the producer organism across runs using varying levels and combinations of sucrose, inulin and tryptone. This utilisation of the Plackett-Burman experimental design allows for the fortification of the growth medium with different levels of these growth factors to optimise the production of bacteriocins by *L. lactis* and determine the optimal combinations that facilitates maximal Nisin synthesis. Each of the 10 runs as outlined in Table 5 was repeated thrice (*n*=3).

**Table 5:** Plackett-Burman Experimental design and coded values to evaluate the effect of variable levels on bacteriocin production by *Lactococcus lactis*.

| Repeat | Variable levels |  |  |  |  | *L. lactis*<br>growth<br>(log<sub>10</sub> CFU/ml) | Bacteriocin<br>production<br>(mg) | Test Organism<br>(log<sub>10</sub> CFU/ml) |
| --- | --- | --- | --- | --- | --- | --- | --- | --- |
|  | Temp | pH | Sucrose | Inulin | Tryptone |  |  | *B. cereus* |
| 1 | -1 | -1 | -1 | -1 | -1 | 7.50 | 65.8 ± 0.012 | 2.80 |
| 2 | -1 | +1 | -1 | 0 | +1 | 8.34 | 72.3 ± 0.015 | 2.42 |
| 3 | -1 | -1 | -1 | +1 | 0 | 8.03 | 74.5 ± 0.011 | 2.17 |
| 4 | -1 | +1 | 0 | -1 | +1 | 8.25 | 70.4 ± 0.018 | 2.22 |
| 5 | +1 | -1 | 0 | 0 | 0 | 7.11 | 68.3 ± 0.013 | 2.40 |
| 6 | +1 | +1 | 0 | +1 | -1 | 8.25 | 71.0 ± 0.017 | 1.94 |
| 7 | +1 | -1 | +1 | -1 | 0 | 8.28 | 75.2 ± 0.014 | 2.44 |
| 8 | +1 | +1 | +1 | 0 | -1 | 8.0 | 73.6 ± 0.016 | 2.83 |
| 9 | -1 | -1 | +1 | +1 | +1 | 8.94 | 75.4 ± 0.019 | 1.95 |
| 10 | -1 | 0 | 0 | 0 | 0 | 7.50 | 70.8 ± 0.010 | 3.64 |

The variable levels coded as -1, 0, and +1 in the table, has the following values: Temp; 30°C (-1) and 37°C (+1); pH; 5.0 (-1), 7.0 (0), and 9.0 (+1); Sucrose; 0 (-1), 15mg/ml (0), and 30mg/ml (+1); Inulin 0% w/v (-1), 1% w/v (0), and 3% w/v (+1); Tryptone 0mg/ml (-1), 5mg/ml (0), and 10mg/ml (+1). Values represent mean ± standard deviation (SD)

These results obtained in this present study are preliminary as other important variables such as aeration and agitation intensity and associated fermenter parameters need to be evaluated before conclusively establishing the levels of fortification to be employed in future works. However, the results obtained from this study can serve as a rough guide for optimal cell growth and bacteriocin production in the interim. Biomass and CFU/ml measurement of bacteria cells were estimated by optical density (OD) measurements and culture (spread plate method). The results show that supplementation with sucrose at 30 mg/ml, inulin at 3% w/v and Tryptone at 10mg/ml gave the highest growth of *Lactococcus lactis* when cultured at a temperature of 30 °C and pH 5.0.

As shown in Figures 7a and 7b, which represent bacteriocin production and bacterial growth, respectively, across the 10 experimental runs (n = 3), the mean bacterial growth was 8.00 ± 0.45 log CFU/ml, while the mean bacteriocin production was 71.49 ± 3.26 AU/ml. The highest bacteriocin yield (75.4 AU/ml) occurred in Run 9, which also recorded the highest growth (8.94 log CFU/ml), indicating a strong positive association between biomass accumulation and bacteriocin synthesis. This trend suggests that optimal metabolic activity in *L. lactis* under favourable nutrient and pH conditions enhances primary metabolite output, consistent with prior findings in lactic acid bacteria fermentations (Akbar et al., 2019, Goyal et al., 2018, Soltani et al., 2022).

**Figure 7.**
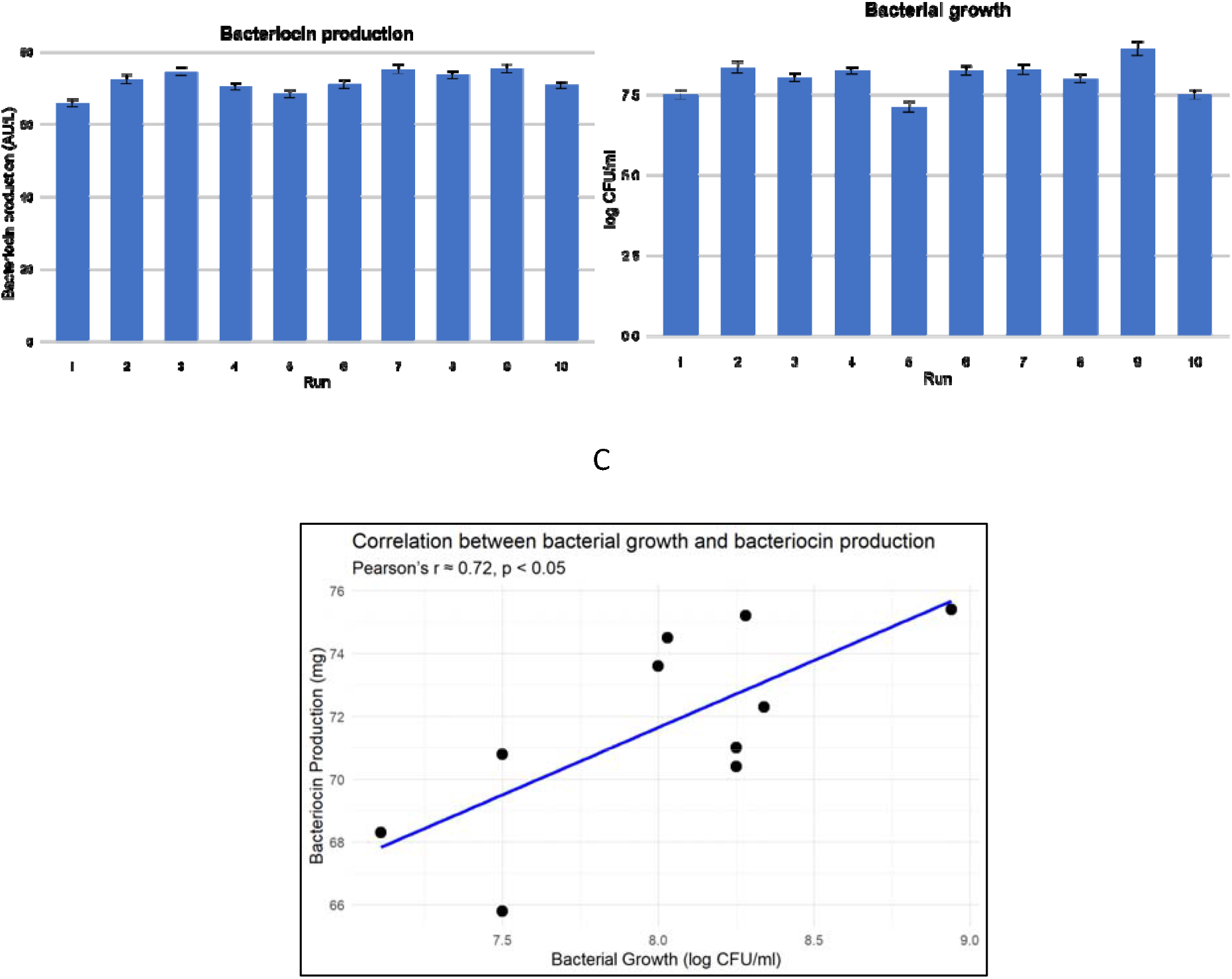
Response results of different variables tested in the PB experimental design on bacteriocin production by *Lactococcus lactis* as a function of a) Bacteriocin production (AU mL); b) Bacterial growth (Log CFU mL^-1^). c) Correlation between bacterial growth and bacteriocin production across all runs. The run number is same as that of the PB experimental design in Table 6. Repeat n=3.

**Table 6.** Effect of temperature and pH alterations on the stability and antimicrobial activity of cell-free extracts of Nisin 2A by *Lactococcus lactis*.

| pH | Zone of inhibition (mm) | Temperature treatments (°C) | Zone of inhibition (mm) |
| --- | --- | --- | --- |
| 4.0 | 15.2 ± 0.014 | 40°C | 16.4 ± 0.012 |
| 6.0 | 16.1 ± 0.012 | 60°C | 16.9 ± 0.014 |
| 8.0 | 15.3 ± 0.015 | 80°C | 16.1 ± 0.011 |
| 10.0 | 14.9 ± 0.013 | 100°C | 16.0 ± 0.013 |

To further explore this relationship, a Pearson correlation analysis was conducted (Appendix 3.4) and shown in Figure 7c. The result showed a strong, statistically significant positive correlation between bacterial growth and bacteriocin production (r = 0.72, p < 0.05), suggesting that increased cell density is closely associated with enhanced bacteriocin yield under the experimental conditions tested. The relatively low variability observed through the error bars in Figure 7a and 7b strengthens the reliability of the strong positive correlation detected between bacterial growth and bacteriocin production.

Interestingly, higher concentrations of the growth factors resulted in the repression of the growth of the lactic acid bacteria, thus these were limited. The application of the Plackett–Burman design in this case was appropriate for the screening and identification of the important factors influencing bacteriocin production in the initial phase. The design provided efficient evaluation of many variables with limited experimental runs. The significant positive correlation between bacterial growth and the amount of bacteriocin produced (r = 0.72, p < 0.05) also indicates the biological reasonableness of the optimisation strategy. These results were statistically verified by replication and correlation analysis, therefore ensuring their validity and justifying their use as a premise for follow-up confirmatory or optimisation studies based on more complex design models such as response surface methodology (RSM).

### 3.6. Biochemical characterisation of Nisin 2A

The biochemical characterisation of Nisin 2A showed a high pH and temperature stability of the peptide. The Nisin 2A fragments were not affected by heat treatment over the temperature ranges of 40-100 °C, retaining its antimicrobial activity, losing only 5% of their activity at 100 °C (Table 6). This indicates the suitability of this peptide for use in food preservation processes that involve high-temperature treatments, such as pasteurization. This result on thermal stability is in agreement with the findings from another high temperature study (Holcapkova et al., 2018) which found that Nisin retained almost 70% of its antimicrobial activity after heat treatment in polylactide at 160 C for 15 minutes. This is similar to a previous report (Akkoc et al., 2011) which highlighted a 50% activity retention after exposure to 121 C for 15 minutes. Similarly, the antimicrobial CFS fractions retained their activity across the pH range of 4.0 to 10.0, losing only 10% of their activity at pH 10 (Table 6), which corroborates the findings of (Goyal et al., 2018). Treatment of the antimicrobial fragments with the enzymes resulted in mixed results. Proteinase K and trypsin resulted in a complete loss in the antimicrobial properties of the extracts, thus confirming the proteinaceous nature of these compounds. However, the extracts were unaffected by catalase while treatment with α-amylase had a slight reduction effect on the antimicrobial activity of the extracts (Figure 8). Enzyme inactivation by Proteinase K and trypsin gives an indication of the proteinaceous nature of the antimicrobial component of the extracts as has been previously observed for most other LAB bacteriocins (Goyal et al., 2018, Furtado et al., 2014). The antimicrobial fragments in the CFS can thus be considered to be a bacteriocin. The biochemical characteristics of cell-free extracts from other isolates in this study is presented with the correlation between bacterial growth and bacteriocin production in Appendix 3.5.

**Figure 8.**
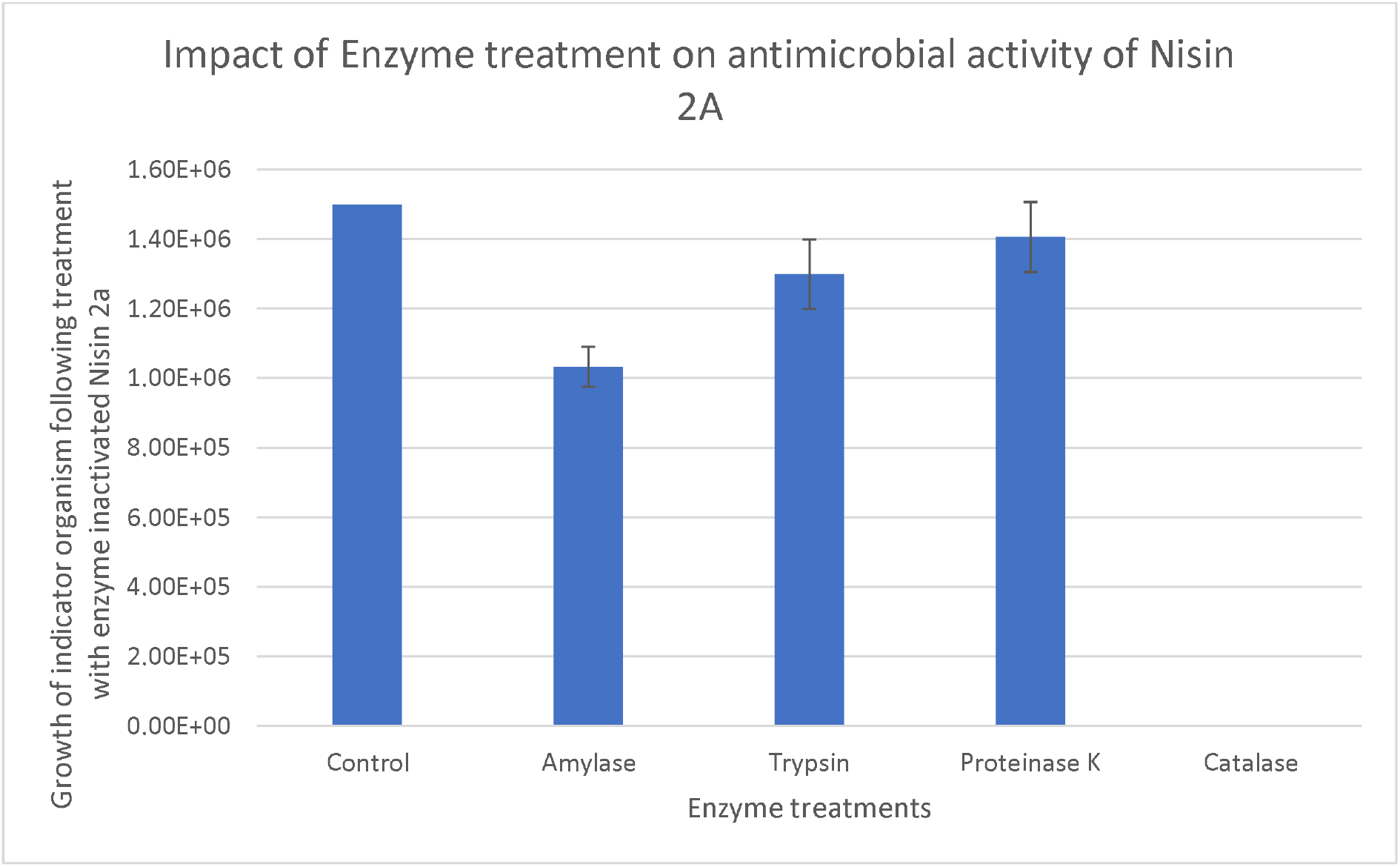
Investigating the susceptibility of Nisin 2A to enzyme treatment/inactivation (n=3) Overall, Nisin 2A from this present study, is stable across a range of environmental treatments, with the resistance of the peptide to high temperatures up to 100°C being of particular significance especially with regards to the control of thermophilic organisms. Furthermore, the degradation of the antimicrobial peptide by Proteinase K and trypsin confirms that the antimicrobial is proteinaceous in nature. Similarly, the slight reduction in the antimicrobial activity of the bacteriocin upon treatment with α-amylase suggests the presence of a carbohydrate moiety within it. It is important to note that the protein degradation observed due to exposure to proteolytic enzymes was similar to results reported by previous studies (Lei et al., 2020, Barman et al., 2018, Ribeiro et al., 2014).

### 3.7. Production and purification of Nisin 2A

Initially following SPE, cation exchange chromatography was performed and 1M NaCl was used to elute the bound Nisin. Obtained product was subjected to SDS-PAGE analysis. However, SDS-PAGE revealed the presence of several other higher molecular weight components. This warranted a further optimisation of the purification process by initiating several trial elution runs using decreasing concentration of NaCl. This was optimised at 400mM as this was shown to be sufficient in eluting the vast majority of the Nisin molecules from the column while keeping the eluant clean/devoid of contaminants as seen in the SDS-PAGE analysis with an approximate size of 3.3kDa (Figure 9), closely aligning with the known molecular mass of Nisin (∼3.4 kDa) as reported by Choi et al. (Choi et al., 2010), and the 3.5 kDa size identified by Abts et al. via tricine-SDS-PAGE (Abts et al., 2011). This finding is significant as it suggests the peptide’s low molecular weight typical of lantibiotics and verifies successful size-based isolation using SDS–PAGE. Quantification using the Pierce protein assay revealed that the total protein concentration of the eluted fraction was 2.9 mg/mL. Further size determination and quantification was achieved using HPLC coupled to a Mass spectrometer.

**Figure 9.**
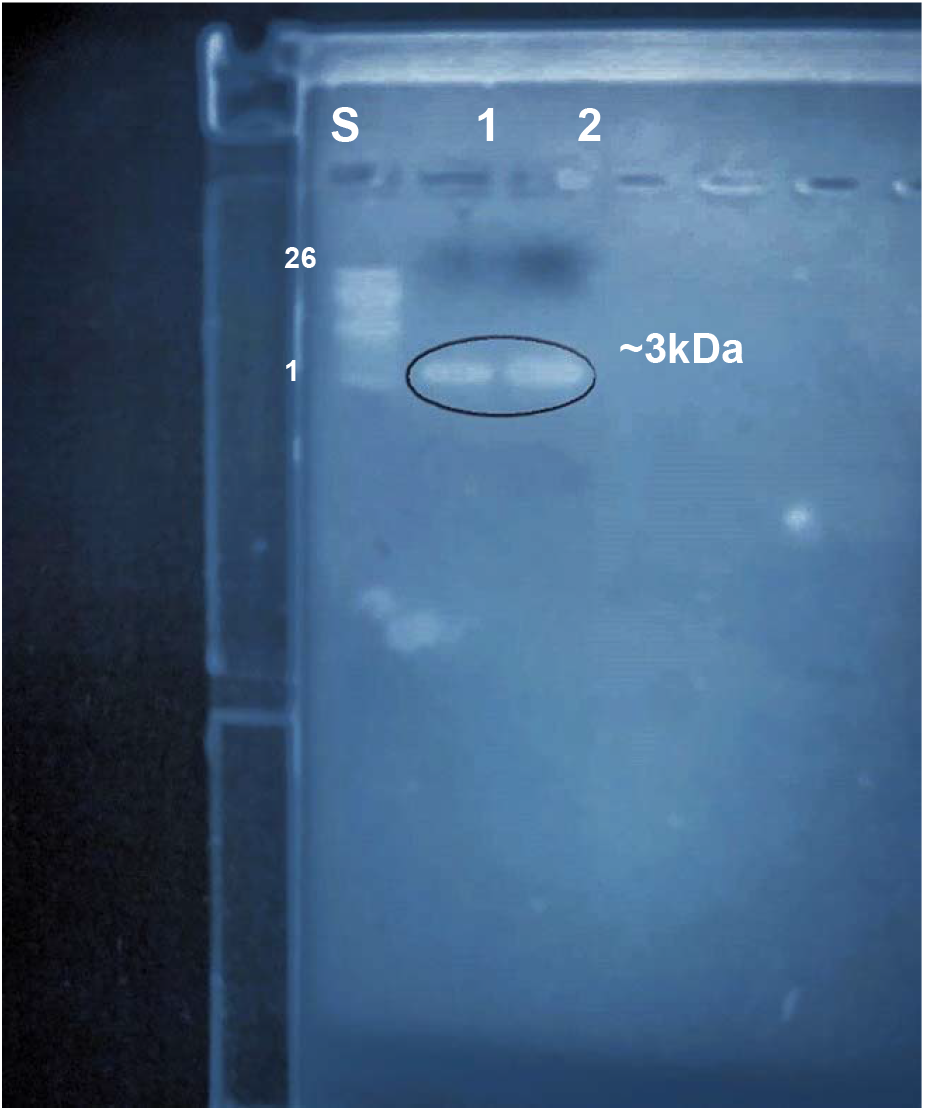
SDS PAGE of Nisin 2A, showing an estimated molecular mass of ∼3 kDa. Lane Standard ladder: 1-26kDa Molecular weight standards (C6210-1VL, Sigma-Aldrich, Gillingham); Lane 1: Nisin 2A; Lane 2: ready-made commercial Nisin (3µg) SDS-PAGE analysis provided a clear separation and visualisation of peptide migration in relation to its molecular mass. However, it is important to consider the limitations of SDS–PAGE for pH-sensitive peptides such as Nisin. Choi et al. (Choi et al., 2010) highlighted that Nisin loses bioactivity at neutral to basic pH levels, including the pH conditions (8.3–8.8) inherent in standard electrophoresis systems. This was not a limitation in this present study as the focus of electrophoresis was a preliminary size determination before mass spectrometry. Also, because of the possibility of losing antimicrobial activity during electrophoresis, the traditional gel-overlay assay post-electrophoresis with inoculated growth medium to study possible zones of inhibition at sites of migration of the bioactive peptide was not undertaken as this was subsequently undertaken in-vitro.

### 3.8. Mass determination of Nisin 2A using HPLC-MS

Following SPE and gel electrophoresis, HPLC-UV analysis was undertaken. HPLC–UV analysis revealed a retention time of 28 minutes for the antimicrobial peptide (Figure 10), indicating successful chromatographic resolution of the target molecule. This retention pattern is consistent with that expected for small, amphiphilic peptides like Nisin, which contain hydrophobic lanthionine and methyllanthionine residues and interact strongly with reversed-phase C18 columns. Moreover, this step added analytical precision to the separation process, allowing partial purification of Nisin 2A prior to biological assays and downstream mass spectrometry.

**Figure 10.**
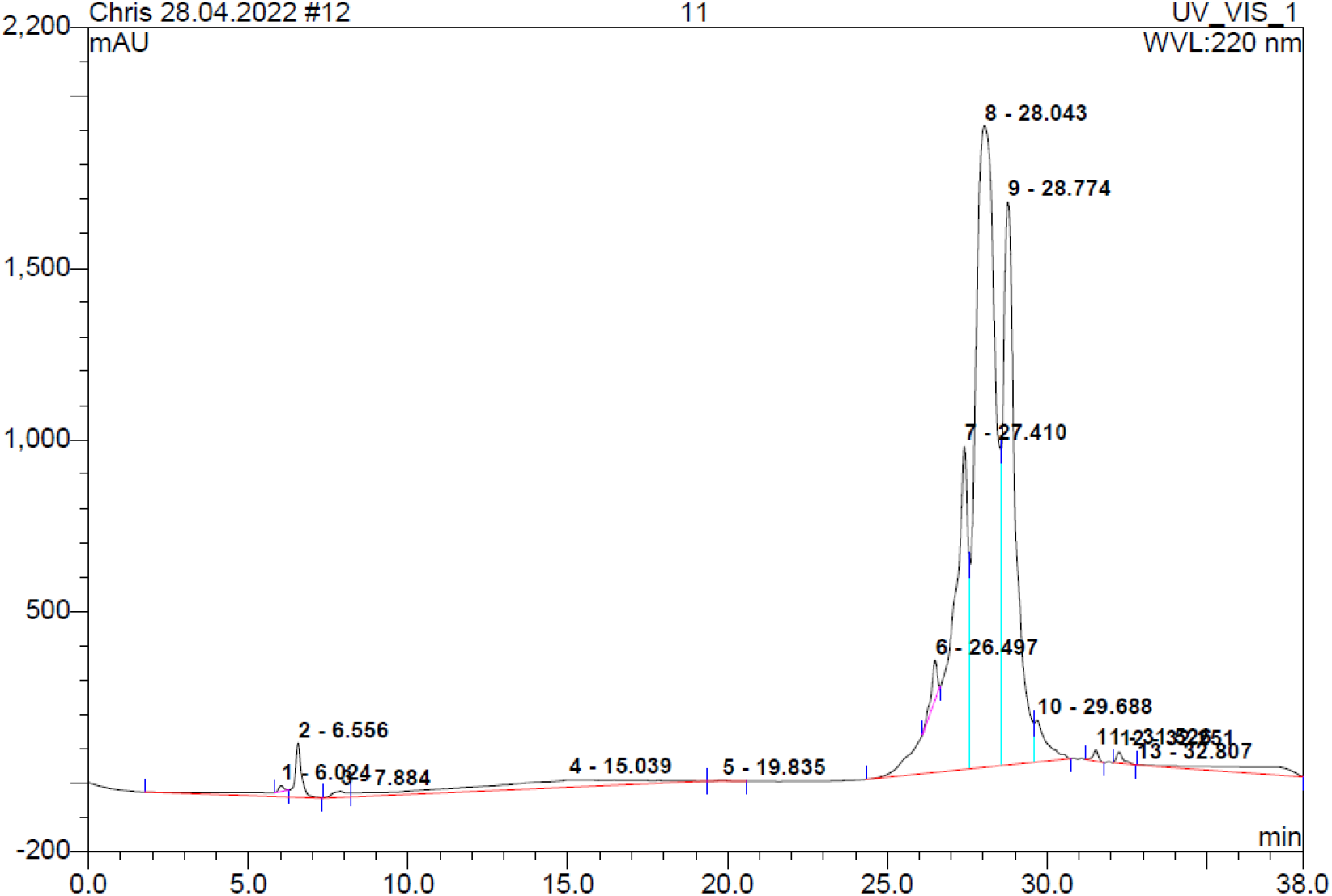
HPLC-UV spectrophotometric separation and detection of Nisin 2A, with peptide elution observed at the 28^th^ minute (peak 8 and 9)

A second separation run was conducted via preparative HPLC to further separate and purify Nisin 2A. The eluants were ultrafiltered using a 1.0 kDa molecular weight cut-off spin filter. UHPLC mass spectrometric analysis yielded only one peptide with a mass fragment of 3,353Da (Figure 11 and Appendix 3.6), which is in agreement with the calculated mass of 3354.07Da for Nisin as outlined in literature (Choi et al., 2010, Abts et al., 2011). Peak integration of the total mass spectrum compared to Nisin standards revealed that the elution fraction contains >98% of Nisin, indicating that this fraction is essentially devoid of contaminants.

**Figure 11.**
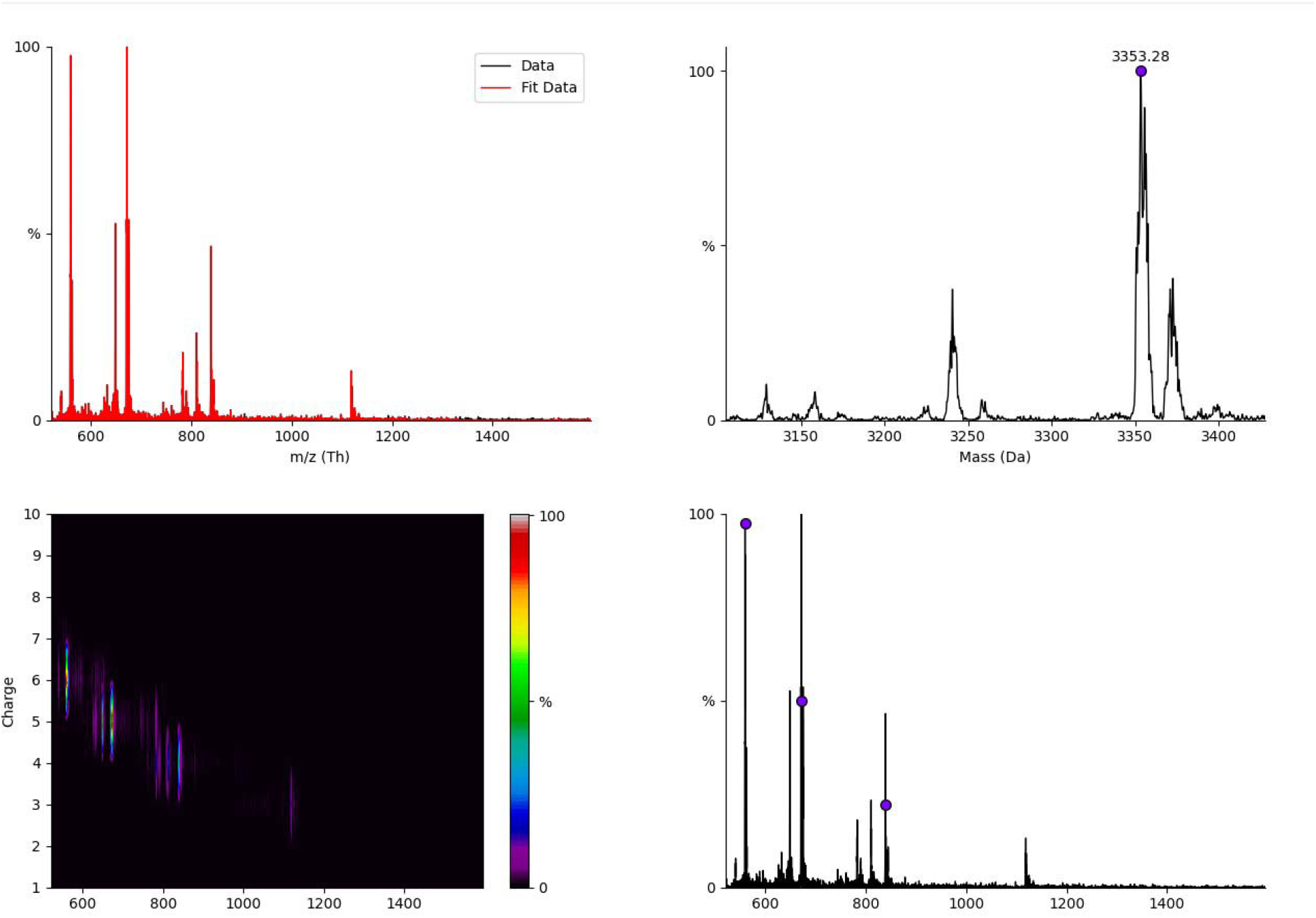
Mass-charge ratio and daughter ion scans (649.1 and 675.3) of purified Nisin 3353.28Da.

A more detailed peptide sequencing, including the use of Nuclear Magnetic Resonance (NMR) is required to conclusively determine if Nisin 2A obtained in this present study is novel and structurally different from previously synthesised Nisin which has been extensively characterised. This is because BLASTP analysis of the genome of the producer *Lactococcus lactis* against known proteins within the prokaryotic antimicrobial peptide database revealed the distinctness of the peptide which although similar to Nisin, did not yield any significant match with any currently known antimicrobial peptide. Although the LC-MS molecular ion peak of Nisin 2A at approximately 3.3 kDa is slightly lower than the standard mass of Nisin A (3.5 kDa), this size variation may not mean a wholly novel peptide, but may indicate structural truncation, post-translational modification, a novel sequence variant or may be due to the impact of the experimental treatments. In contrast, Ko et al. (Ko et al., 2015) identified Nisin A and Z with parent ions at 671.6 and 667.0 m/z respectively, and corresponding fragment ions, confirming the known structures of these lantibiotics. In the current study, the detection of Nisin 2A by LC-MS at a similar mass-to-charge ratio of 649.1 and 675.3 supports its classification within the Nisin family, while suggesting it is structurally distinct. These results indicate the identification of a potentially novel antimicrobial compound that may offer more functionalities or target a broader spectrum of pathogens compared to the standard Nisin. While microbes have not developed any notable spontaneous resistance to known variants of Nisin, it has been documented that certain bacterial strains exhibit intrinsic resistance to Nisin through a variety of mechanisms, including the production of resistance proteins, biofilm development, and cell wall alteration (Draper et al., 2015, Field et al., 2019). Hence, this peptide if proven novel could serve as an alternative or complementary antimicrobial agent in the food industry, especially in systems where the efficacy of Nisin may be limited by microbial resistance.

## CONCLUSION

This study presents the successful isolation, screening, molecular characterization, and functional analysis of a novel bacteriocin-producing strain of *Lactococcus lactis* designated *L. lactis* Lac001 from brined cheese. Among five lactic acid bacteria (LAB) isolates tested, *L. lactis* exhibited the highest antimicrobial activity against *Bacillus cereus*, a significant foodborne pathogen, confirming its suitability as a source of potent antimicrobial peptides. In silico screening using antiSMASH 7.1.0 detected multiple biosynthetic gene clusters (BGCs), including those responsible for producing lanthipeptides (Class I), T3PKS, betalactones and RiPP-like clusters, with similarities to known Nisin and lactococcin operons. These features underscore the strain’s genetic potential for antimicrobial compound biosynthesis. Despite the inability to conclusively map the gene cluster for Nisin 2A, predicted BGCs demonstrated high similarity to Nisin A gene clusters and lactococcin BGCs, suggesting evolutionary linkage. Also, through the predicted structure and molecular weight of Nisin 2A and with BLASTP analysis showing no exact matches to known bacteriocins, this strongly suggest that Nisin 2A may be a previously undescribed peptide with novel sequence or post-translational modifications. Cultivation of *Lactococcus lactis* and bacteriocin synthesis was achieved using modified MRS medium. The Plackett-Burman experimental design was used to optimize bacteriocin yield and identify sucrose, inulin, and tryptone as optimal nutritional supplements and key enhancers of both biomass and antimicrobial production, with a significant correlation between biomass accumulation and bacteriocin activity *p* < 0.05, reinforcing the close metabolic link between growth conditions and metabolite synthesis. In this study, obtained bacteriocin was purified using a multi-step protocol involving ammonium sulphate precipitation, SPE, and HPLC, and its mass was validated through UHPLC-MS and SDS-PAGE analysis. Comprehensive biochemical, genomic, and proteomic analyses confirmed that *L. lactis* Lac001 produces a distinct bacteriocin in the Nisin family, tentatively named Nisin 2A, with a molecular mass of approximately 3.3 kDa as determined by UHPLC-MS and SDS-PAGE. This places it within the size range of Class I lantibiotics but slightly below the size of known Nisin variants such as Nisin A (∼3.5 kDa), suggesting it may represent a distinct structural variant or truncated form with unique functional properties. The purified peptide demonstrated broad-spectrum antimicrobial activity, particularly against the test Gram-positive bacteria *Bacillus cereus* and retained its bioactivity across a wide pH range (3–9) and high thermal conditions (up to 100 °C), indicating high physicochemical stability, traits advantageous for food preservation. Furthermore, it had high sensitivity to proteolytic enzymes (Proteinase K and Trypsin), confirming its proteinaceous identity. Notably, the peptide was thermostable and retained up to 90% of its initial activity after thermal treatment, and maintained consistent inhibitory performance after extended storage, further highlighting its application potential in real-world food systems. Its small molecular size may confer improved diffusibility and membrane-targeting efficiency, making it an ideal candidate for use as a natural bio-preservative in minimally processed foods, particularly where thermal stability and broad antimicrobial coverage are required. Further studies including a more detailed structural elucidation, study of bacteriocin mode-of-action, microencapsulation and food matrix application trials will aid to fully exploit its potential in the food and pharmaceutical sectors.

